# Polyphyly of the *Niphargus stygius* species group (Crustacea, Amphipoda, Niphargidae) in the Southern Limestone Alps

**DOI:** 10.1101/2022.04.28.489871

**Authors:** Fabio Stoch, Alice Salussolia, Jean-François Flot

## Abstract

The *Niphargus stygius* species complex is a groundwater group of large-sized, sexually dimorphic species inhabiting mainly caves and, less frequently, wells and springs. According to the taxonomists of the last century, this species complex was supposed to be present in the whole Southern Limestone Alps of Italy as well as in peninsular Italy, Slovenia, and Croatia. Considering the large, presumed distribution area, we tested the contrasting hypotheses of monophyly versus paraphyly of this subterranean species complex, taking in account the presence of putative cryptic species. For this reason, we sampled the type localities of all the described species in the complex present in the Italian Southern Limestone Alps and neighbouring areas, and used nuclear rDNA (28S, ITS region) and mtDNA (COI) markers to assess the phylogenetic relationships and species richness. Phylogenetic analysis confirmed that the *Niphargus stygius* complex in the Southern Limestone Alps is polyphyletic comprising an eastern clade (the *N. julius* clade, present in NW Italy, northern Slovenia, and southern Austria) and two western clades (the *N. brixianus* and *N. montellianus* clades). These two clades are not closely related to the eastern one but rather form a monophyletic group together with a widely distributed Apennine clade (*N. speziae* clade). None of these clades is closely related to typical *N. stygius*. Three different molecular species delimitation methods applied to COI and rDNA sequences recognized slighlty different numbers of putative species, suggesting that each clade is a species complex. Bayesian time-calibrated phylogeny revealed that most clades began to split up during Miocene and Pliocene, ruling out the effect of Pleistocene glaciations, evidenced only by the COI marker, in explaining their speciation process and justifying the presence of several putative cryptic or pseudocryptic species.

## Introduction

With more than 425 species described (Horton et al., 2021), Niphargidae are the most species-rich taxon of freshwater amphipods in the world (Väinölä et al., 2008). The family Niphargidae is distributed in the Western Palearctic (Borko et al., 2021), including most of Europe, mainly – but not exclusively – in groundwaters south of the Pleistocene ice sheet boundary (McInerney et al. 2013; Zagmajster et al., 2014). The genus *Niphargus* was described by Schiödte (1849), and *Niphargus stygius* (Schiödte, 1847) was designated as its type species (type locality: Postojna cave in Slovenia). The taxonomists of the last century considered this species as quite widespread. In the pioneering paper by D’Ancona (1942), most of the known species and subspecies described during the first half of the last century for the genus *Niphargus* were considered as synonyms of *N. stygius*. However, despite his exhaustive morphological study of minute morphocharacters, the conclusions of D’Ancona’s (1942) monograph were considered erroneous in the following years when taxonomists identified several distinct morphotypes within the genus *Niphargus* and assigned species to subgenera or ‘species groups’ (Karaman S., 1952; Straškraba 1972). Unfortunately, the high number of species, the unknown variability of the morphocharacters used for species identification, and the varying quality of their descriptions scattered over several local journals (Fišer et al., 2009a), i.e., in ‘grey literature’, prevented taxonomists from achieving satisfactory results.

The most important contribution to the assessment of ‘species groups’ within niphargids was the one by Straškraba (1972), reported in the paper concluding the proceedings of the “1er Colloque International sur le genre *Niphargus*” held in Verona in 1972. In this article, the genus *Niphargus* s. str. was divided into 13 ‘species groups’ plus a fourteenth group comprising 10 species ‘incertae sedis’. The ‘*stygius*-group’ encompassed several species from Slovenia and northeastern Italy, while the Apennine species were allocated, following the suggestion of another contribution published in the same proceedings (Vigna Taglianti, 1972), in the ‘*speziae-romuleus-tatrensis’* group, including also the *Niphargus tatrensis* species complex distributed in central and eastern Europe. Later, following the recommendations of Straškraba (1972), *N. stygius* was redescribed by Sket (1974) as the representative of its own ‘group’. This picture was turned over by G. Karaman (1993), who, in his monograph on the fauna of Italy, reassigned all the different species and subspecies of the *‘speziae-romuleus’* group to the single species *Niphargus stygius*. These conclusions were criticized by Stoch (1998), who considered the species present in the Southern Limestone Alps in Italy as different, albeit belonging to the well-established *‘N. stygius* group’.

Recently, molecular and morphological systematic studies suggested that the classification proposed by Straškraba (1972) was not justified from a phylogenetic perspective (Trontelj et al. 2009; Fišer et al. 2009b). As regards *N. stygius*, Delić et al. (2017) used molecular taxonomy to revise the populations present in Slovenia and northwestern Croatia where S. Karaman (1952) had subdivided *N. stygius* into seven subspecies. Using uni- and multilocus delineation methods, Delić et al. (2017) showed that the group in the northern Balkan area was not monophyletic but consisted of 15 parapatric and sympatric cryptic species, which were described using molecular diagnoses. Later, Stoch et al. (2020) revised the *N. tatrensis* species complex, demonstrating that there was no relationship between this old, monophyletic lineage and neither typical *N. stygius* nor *N. speziae* and *N. romuleus* from the Apennines, thereby rejecting the hypothesis of Vigna Taglianti (1972). Finally, *Niphargus stygius* itself was considered as a species complex (Delić et al., 2021), and at least four different lineages were recognized even within the small distribution area of this species encompassing the karstic areas of northeastern Italy and western Slovenia. No other paper has addressed the molecular taxonomy of Italian pre-Alpine species attributed to the *Niphargus stygius* group so far.

The lack of phylogenetic knowledge for wide areas as regards groundwater niphargids and the presence of cryptic and pseudocryptic species are the worst-case scenario of morphotaxonomy’s inadequacy to describe biodiversity. This is not only an exclusively taxonomic problem; no ecological information on these cryptic species is in most cases available, raising conservation issues (Delić et al., 2017).

Given these premises, we present herein a molecular phylogeny of the various species inhabiting the Southern Limestone Alps in Italy and neighbouring areas of Austria and Slovenia, traditionally attributed to the *Niphargus stygius* species complex (Stoch, 1998). The studied group includes morphologically similar populations distributed from Lake Como along the Southern Limestone Alpine chain to western Slovenia and southern Austria. Our study has three major aims. First, to reconstruct the phylogeny of this ‘western species group’ of the *Niphargus stygius* complex so that we can test the hypothesis that it constitutes a monophyletic evolutionary unit and assess its relationships with *N. stygius* as well as morphotaxonomically similar clades. Second, to test the conjecture that the species complex comprises multiple cryptic species in this study area and apply molecular species delimitation methods to identify them. Third, to reconstruct a time-calibrated phylogeny to shed light on the origin and separation of the different species in the area.

## Material and methods

### Sampling design

The Southern Limestone Alps (or Prealps) are the offshots of the Eastern Alps south of the Central-Eastern Alps. Their greatest extent is in Italy, while extending eastwards in the nearby Austria and Slovenia. The Southern Limestone Alps extend from the Como Lake valley in Lombardy to the Pohorje in Slovenia. They are characterized by a West-East succession of highly karstified carbonate massifs, separated by deep valleys (in the eastern part occupied by lakes of glacial origin, the larger being the lakes Como, Iseo, and Garda).

Samples were collected prospecting 129 sites (caves, wells, and springs) throughout the wide range of the so-called *Niphargus stygius* species group as defined by Karaman (1993) and Stoch (1998) in Italy and southern Austria (Fig. 1 and Tab. S1). Many species of this group occur at very low densities exclusively in caves, where sampling of groundwater fauna is logistically demanding (Fišer & Zagmajster, 2009; Pipan & Culver, 2007), requiring a proper equipment and a good knowledge of progression in cave meanders and wells using ropes and specialized techniques. All the type localities of the different species and subspecies of the previously defined *N. stygius* group described from Italy were sampled to build a complete reference library.

**Figure 1.**
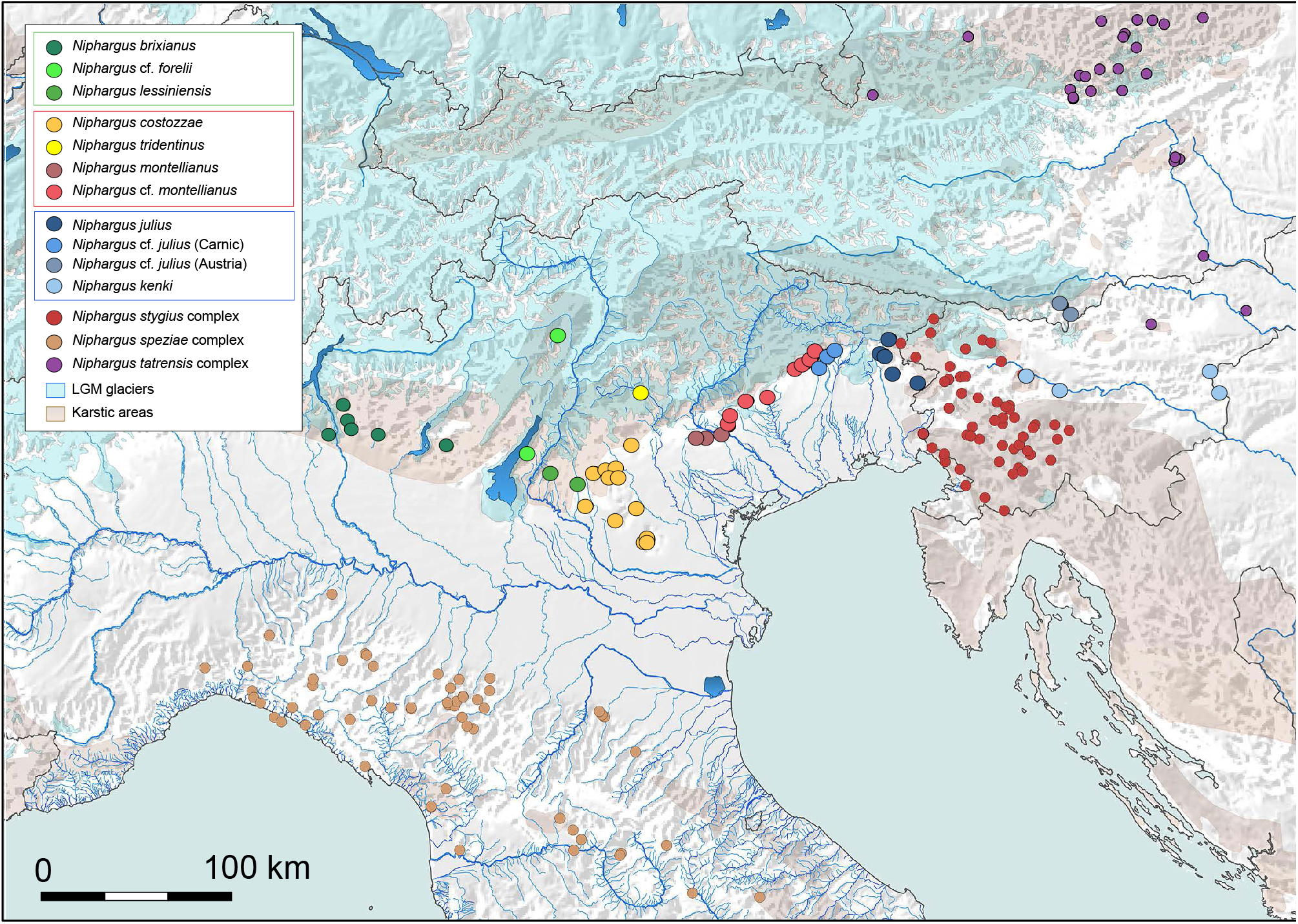
Map of the study area showing the distribution of the sampling sites of specimens of the *Niphargus stygius* group; species names are based on morphology and follow mainly Stoch (1998). The extent of karstic areas and of the glaciers during the Last Glacial Maximum (LGM) are reported as well.

Cave-dwelling specimens were collected by one of the authors (F.S.), as well as by several speleologists (mentioned in the acknowledgements). In caves, most of the species are present in pools fed by percolating waters as well as in small siphons and subterranean lakes; in some cases (like Grotta della Bigonda, Trentino) the access to the cave needed to empty the entrance siphon using an electric pump. In few cases, specimens were present in deep lakes in quite complex karstic systems reaching the saturated zone and were collected by speleodivers. Sampling was carried out using hand nets, and by positioning baited traps (bottles containing chicken liver) in siphons and small lakes overnight. Spring specimens were hand-netted, or drift nets were positioned for a few hours at the spring outlet. Finally, artificial wells were sampled either using Cvetkov nets (Cvetkov, 1968) or by positioning baited traps at the bottom of the wells. The samples are deposited in the collection of the Evolutionary Biology & Ecology unit of the Université libre de Bruxelles (ULB), Belgium, stored in freezers at -20°C. Samples used for morphotaxonomical analyses are stored in the first author’s personal collection in Rome and in the Natural History Museum of Verona, Italy. All the information on the specimens used in the analyses, georeferenced collecting sites and DNA vouchers are available in Supplementary Material, Table S1.

Additional sequences were downloaded from GenBank, where results of previous studies on the *Niphargus stygius* complex in Slovenia and Croatia (Delić et al. 2017, 2021) and of the alleged similar *Niphargus tatrensis* complex in Austria and Eastern European countries (Stoch et al. 2020) are deposited. Moreover, all the sequences reported in Borko et al. (2021, supplementary material) obtained from 385 niphargid and pseudoniphargid species, complemented by those reported by Weber et al. (2021), were downloaded.

### Phylogenetic and species delimitation analyses

One or two pereopods of each of the 75 selected specimens (see Tabs. S1 and S2) were used for DNA extraction, and the remaining parts of each specimen were stored in 96% EtOH at -20°C at ULB. Extraction of genomic DNA was performed using the NucleoSpin@ Tissue kit by Macherey-Nagel, following the manufacturer’s protocol. The eluted DNA was stored at 4°C until amplification, then long-term stored at -20°C. The following markers were PCR-amplified: (1) a fragment (between 987 and 993 bp long) of the nuclear 28S rRNA gene (Verovnik’s fragment: external primers of Verovnik et al. 2005 complemented with internal primers of Flot et al. 2010) ; (ii) a 658 bp fragment (Folmer’s fragment: Folmer et al. 1994, using the primers of Astrin & Stüben 2008) of the mitochondrial cytochrome *c* oxidase subunit I (COI); (3) the complete internal transcribed spacer (ITS) region (together with flanking portions of the 18S and 28S genes and including 5.8S) using the six primers described in Flot et al. (2010). A list of primers and PCR amplification protocols used in this study is available in Tab. S3. Direct sequencing was performed using the same primers as for amplification as well as internal primers (Flot et al. 2010); 28S and COI PCR products were sent for bidirectional Sanger sequencing to Genoscreen (Lille, France), while ITS products were sent to Macrogen Europe (Amsterdam, The Netherlands).

Chromatograms were inspected, assembled, and cleaned using the program Sequencher 5.4.6 (Gene Codes). Some 28S and ITS chromatograms contained double peaks, as expected in the case of length-variant heterozygotes (Flot et al. 2006); these individuals were phased using the web tool Champuru (Flot 2007, available online at https://eeg-ebe.github.io/Champuru). The two alleles of the same heterozygous individuals are indicated with the letters ‘a’ and ‘b’ following the voucher code (Tab. S1).

The position of the *Niphargus stygius* -like species complexes of the Southern Limestone Alps and of the other species groups cited in the taxonomic literature as related to these populations (E-Slovenian and Croatian *Niphargus stygius* group. *N. speziae* group and *N. tatrensis* group) within the phylogenetic tree of Niphargidae were inferred by comparison with 183 other niphargid species (Tab. S2) using as outgroups the family Pseudoniphargidae (genera *Microniphargus* and *Pseudoniphargus*), which was assessed as the sister group of Niphargidae in our previous study (Weber et al., 2021). For this phylogenetic analysis, we screened the molecular dataset assembled by Borko et al. (2021), including 385 species. Considering that the dataset included several species represented only by mitochondrial COI partial sequences and most of their markers were not used in our analyses, we revised and pruned the dataset as follows: we retained only complete (or almost complete) sequences (when these sequences were related to those of the target clades, we performed the phylogenetic analyses before and after removing them, to check for any possible loss of information); likewise, we retained only the markers we used in our analyses (COI, ITS region, and 28S-I fragment), while short fragments of further markers, i.e. histone H3, 18S rRNA and another 28S fragment, as well as partial sequences of the phosphoenolpyruvate carboxykinase (PEPCK), glutamyl-prolyl-tRNA synthetase (EPRS), heat shock protein 70 (HSP70), and arginine kinase (ArgKin) were not used, considering they did not change the results of our analysis. Some missing ITS sequences were added using our own data from specimens of the same locality. All in all, 253 sequences were retained in the analysis (including our newly obtained sequences -see Tabs. S1 and S2), including as outgroup three species of Pseudoniphargidae. Considering that COI marker can show saturation in such an old family group, we tested for codons saturation plotting observed pairwise divergences for both transitions and transversions against genetic distance (p-values) using DAMBE 7 (Xia and Lemey, 2009) and calculated an index of saturation for every codon following Xia et al. (2003). The results, shown in Fig. S1, demonstrated that the first two COI codons did not show any sign of saturation, while the third codon clearly showed a large dispersion in transversion and a slight saturation in transitions. Albeit saturation for the third codon was not statistically significant (being the observed Iss lower than IssSym assuming bothan asymmetricl than a symmetrical topology), the graphs clearly demonstrated that the third codon could influence the whole pattern and branch support. For this reason, only the first and second COI codons were used to build the global niphargid trees.

All sequences were aligned for each marker using the E-INS-i algorithm implemented in MAFFT 7 (Katoh and Standley 2013), and the optimal substitution model was selected for each marker using ModelFinder (Kalyaanamoorthy et al. 2017, implemented in the IQ-TREE 2 software package) according to the Bayesian Information Criterion (Schwarz 1978). Phylogenetic trees were obtained for both the ITS-28S region and the COI region to check for mitonuclear discordance. Considering that no significant discordance was detected, phylogenetic analyses were performed by concatenating the aligned rDNA long fragment with COI Folmer’s fragment 1st and 2nd codons, while the full COI fragment was used for a more detailed tree built only for the target *Niphargus stygius* group. Phylogenetic relationships were reconstructed on the concatenated dataset using maximum likelihood and 1,000 ultrafast bootstrap replicates (Minh et al., 2013) in IQ-TREE 2 (Minh et al. 2020); two partitions were used, the first one for nuclear rDNA markers (substitution model GRT+G+I) and COI (substitution model TIM+F+I+G4).

After the ML phylogenetic analysis, a time-calibrated Bayesian phylogeny was reconstructed on the same concatenated dataset in BEAST (Bayesian Evolutionary Analysis Sampling Trees) 2.6.1 (Bouckaert et al. 2019), following the best-fit model of evolution proposed by the bModelTest (Bouckaert and Drummond 2017) package. Substitution models and clock models were unlinked for the two partitions (ITS region + 28S and COI). Based on marginal likelihood (Path Sampler extension: Baele et al. 2016), a Yule speciation tree prior was used for the analyses. To account for lineage-specific rate heterogeneity, a lognormal relaxed clock (Drummond et al. 2006) was used. The coefficient of variation (CV) reported in Tracer 1.7 (Rambaut et al. 2018) employing the relaxed clock was higher than 0.1, suggesting that this clock fitted better the dataset than a strict clock (Drummond and Bouckaert 2015). Unfortunately, no fossil is known in the *Niphargus stygius* complex; for this reason, we used the calibration points as in Borko et al. (2021). Four independent runs of 100,000,000 generations sampled every 1,000 steps were performed and combined using LogCombiner 2.6.1 included in the BEAST package. The stationarity of each single run was checked in Tracer 1.7 (Rambaut et al. 2018). The first 10% of the trees were discarded as burn-in and the remaining samples from the posterior distribution were summarized using TreeAnnotator (included in the BEAST package) in the maximum clade credibility tree.

Phylogenetic networks were built for both mtDNA (COI) and nuclear rDNA (ITS+28S-I) sequences using HaplowebMaker (Spöri & Flot 2020, available online at https://eeg-ebe.github.io/HaplowebMaker/).

The following molecular species delimitation methods (SDMs) were applied to both mtDNA and rDNA markers: (i) a distance-based method, i.e. ASAP (Assemble Species by Automatic Partitioning: Puillandre et al. 2021: https://bioinfo.mnhn.fr/abi/public/asap/asapweb.html) using p-distances (p<0.05), ; (ii) a tree-based method, i.e. the most recent version of PTP (Poisson Tree Processes: Kapli et al. 2017) using the webserver https://mptp.h-its.org/#/tree at p<0.001; (iii) a fixed threshold method (TH) implemented by Lefébure et al. (2006) based on the observation that two clades diverging by more than 0.16 substitution per site, as measured by patristic distances along a COI ML likelihood tree, have a strong probability (p<0.01) of belonging to different morphospecies. Albeit haplowebs were built following Spöri & Flot (2020), they were not used for species delimitation considering the very low amount of heterozygote individuals for the nuclear rDNA region present in the dataset.

## Results

### Phylogenetic analysis

The phylogenetic analysis demonstrated that Straškraba’s (1972) *Niphargus stygius* group in Italy and bordering regions (Fig. 1) is polyphyletic (Fig. S2). The detailed ML tree reported in Fig. 2, showed that at least two large unrelated clades can be recognized in the Italian Pre-Alps, and neither of the two is a sister group of *Niphargus stygius* or any of the other members of the *N. stygius* species group. The strongly supported clades (Fig. 2, 100% bootstrap support) were: (i) the *Niphargus julius* clade, distributed in the Eastern Alps in Italy, Austria, and Slovenia (Fig. 1) surrounding the areal of *Niphargus stygius*; (ii) a widely distributed Alpine-Apennine clade (Fig. 1). This large clade included: (i) the *Niphargus brixianus* (sub)clade, distributed in the western part of the Southern Limestone Alps (Lombardy and Veneto/Trentino including the western Lessinian massif); (ii) the *Niphargus costozzae* (sub)clade, distributed in the eastern part of the limestone Venetian Pre-Alps; (iii) the *Niphargus montellianus* (sub)clade, present in the conglomeratic massifs of eastern Venetian Pre-Alps and in limestone massifs of Carnic Pre-Alps; (iv) the *Niphargus speziae* clade, present in the Apennines. This last clade will not be considered further in the present paper, being absent from the study area.

**Figure 2.**
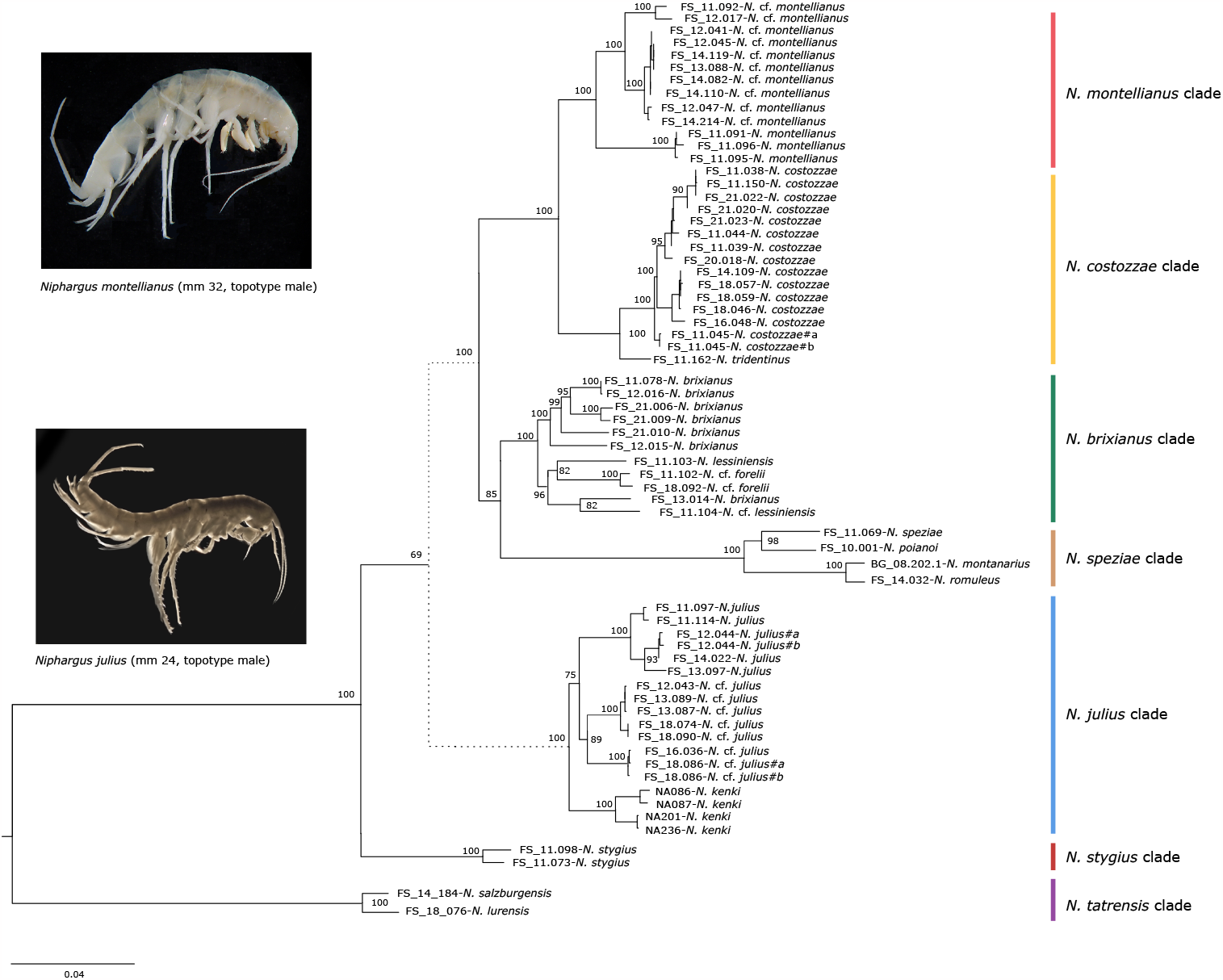
ML phylogenetic tree of the *Niphargus stygius* group using a concatenated set of markers (ITS region, 28S-I and COI). Other clades were used as outgroups following the results of previous analyses (see Tab S1). Bootstrap values of very recent splits, that overlapped with species labels, were omitted for clarity (all either 99 or 100%)

### Molecular species delimitation methods

Results of SDMs are reported in Tab. S4; additionally, ML phylogenetic trees used for PTP are reported in Fig. S3.

Western pre-Alpine clades (*N. brixianus, N. costozzae*, and *N. montellianus* clade: Tab. S4): ASAP and PTP applied to nuclear rDNA marker (ITS+28S-I) gave a very conservative putative species estimate (4), splitting *N. montellianus* into two subclades (Carnic and Venetian pre-Alpine ones: Fig. 3a). Whereas, applying these methods to COI yielded a higher putative species number (15 for ASAP and 14 for PTP), splitting especially the *N. brixianus* clade (Fig. 4a). The threshold method for COI recognized 7 putative species (Tab. S4).

**Figure 3.**
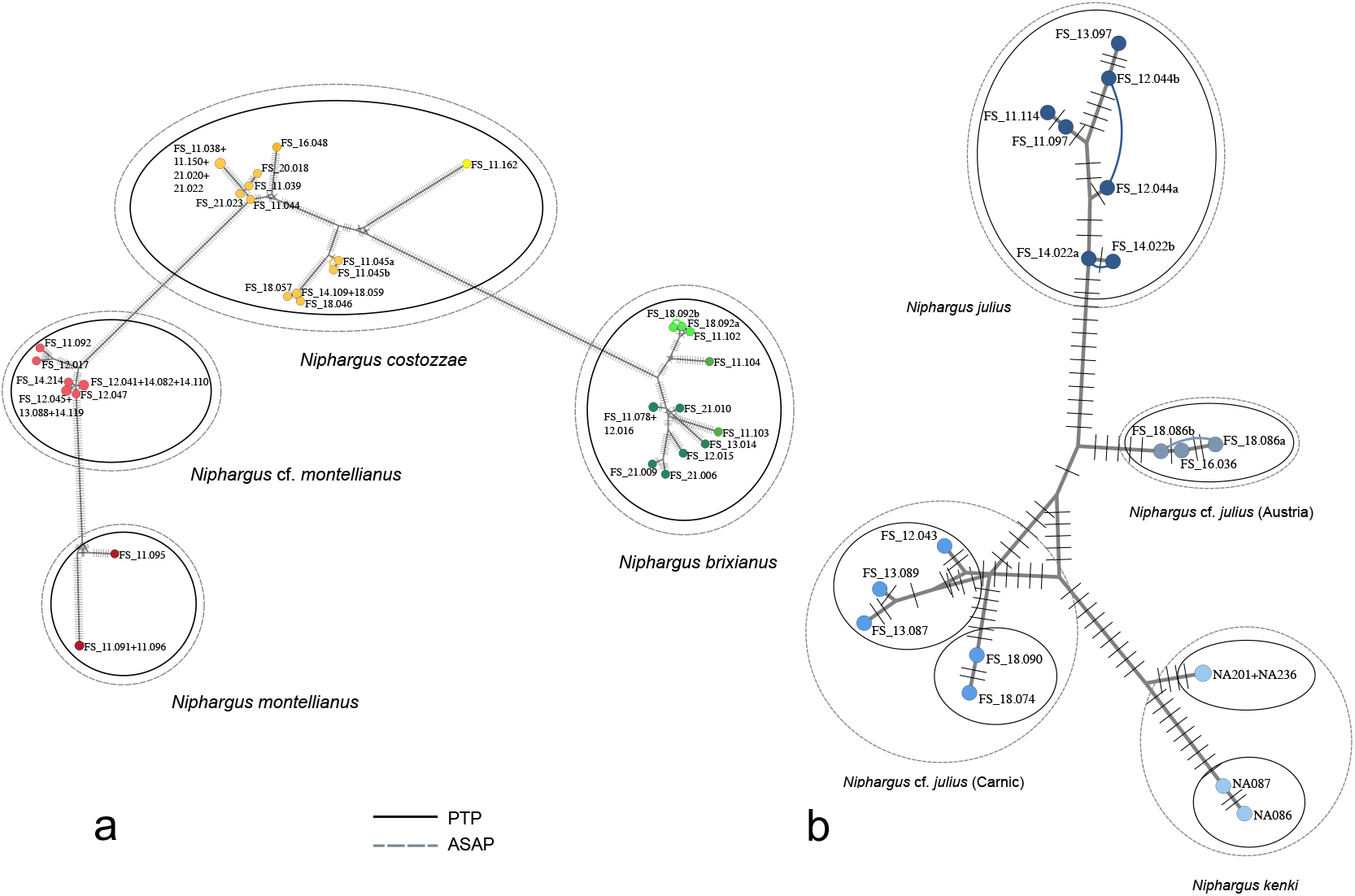
Median joining network of the ITS+28S-I (nuclear rDNA) sequences of the of the *Niphargus stygius* group obtained in the present study. Alleles of the same indivilual are connected by an arc, turnint the haplotype network in a haploweb.

**Figure 4.**
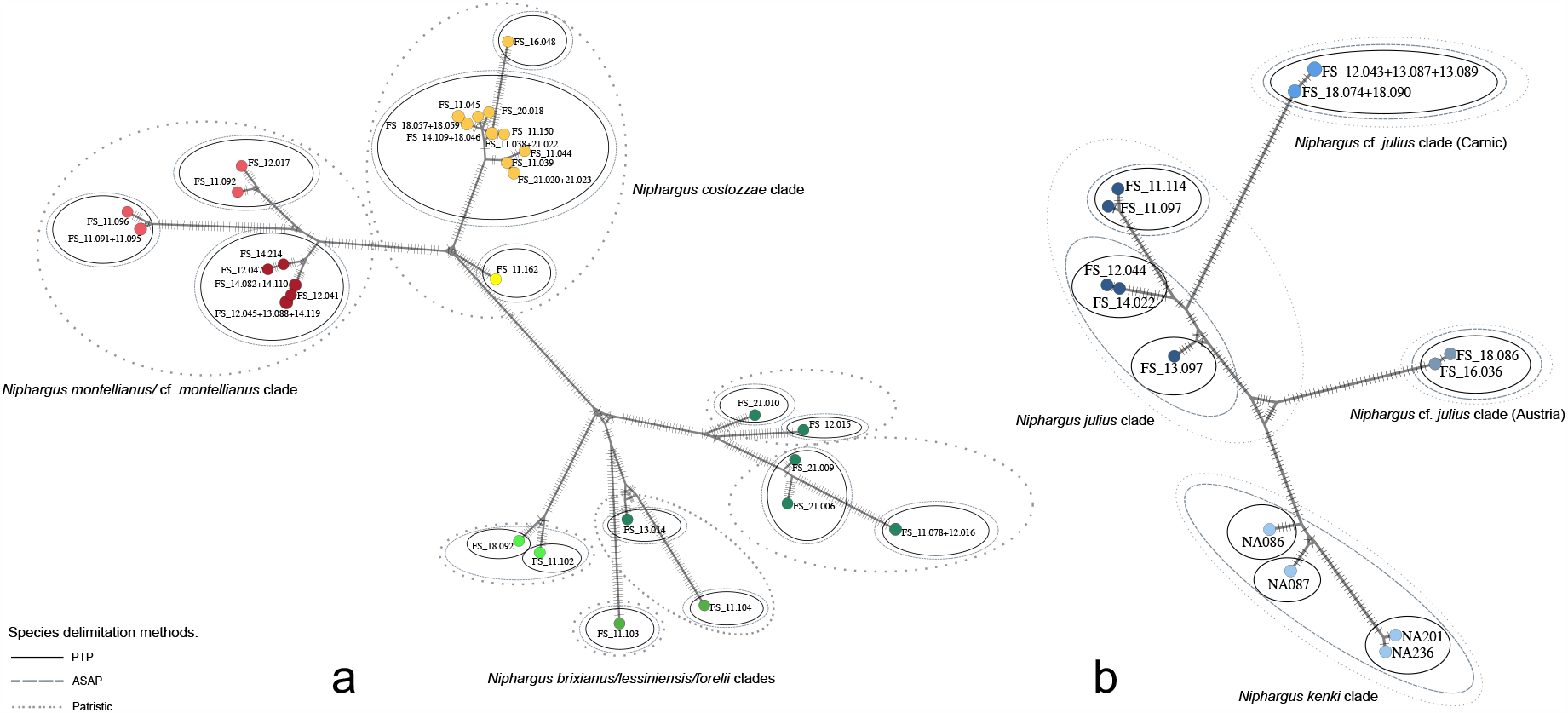
Median joining network of the COI (mt DNA) sequences of the *Niphargus stygius* group obtained in the present study.

Eastern clade (*N. julius* clade: Tab. S4): ASAP and PTP applied to ITS+28S-I (Fig. 3b) recognized respectively 4 (Carnic, Julian, Slovenian, and Austrian) and 6 putative species (splitting both the Carnic and the Slovenian clades in two putative species each). The application of these methods to COI gave again a slightly higher putative species number (5 for ASAP and 8 for PTP) (Fig. 4b). The threshold method for COI recognized 4 putative species, coinciding with ASAP results applied to rDNA markers.

### Time-calibrated phylogeny

The polyphyly of the *Niphargus stygius* group is well illustrated by the time-calibrated phylogeny obtained using the Bayesian analysis (Fig. 5). Results were very similar to those reported by the ML phylogenetic tree. The temptative calibration points applied to the timetree suggested that the most ancient splits between the main recognized clades in the Western Southern Limestone Alps took place in the Miocene, i.e., around 8-10 Ma, while spilts inside the eastern *N. julius* clade were more recent. The separation of most of the minor clades identified in the tree took place during the Pliocene and then followed the vicissitudes of the Pleistocene glaciations.

**Figure 5.**
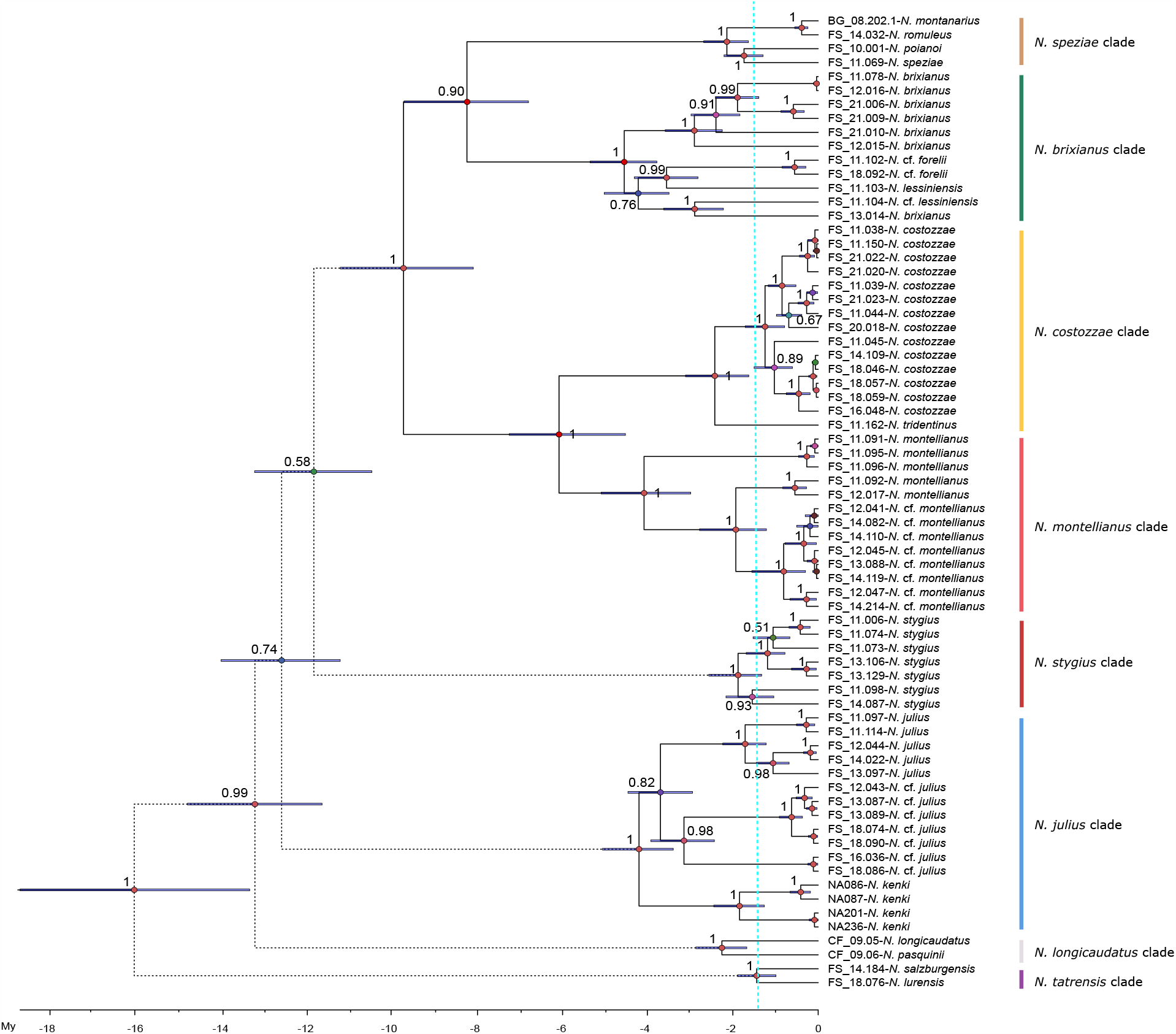
Time-calibrated maximum clade credibility tree of the *Niphargus stygius* group derived from a BEAST analysis of concatenated ITS, 28S-I and COI sequences (1st and 2nd codon). Both the posterior probabilities of the nodes and the 95% confidence intervals of their ages are reported.

## Discussion

### The *Niphargus stygius* group in Italy

The results illustrated above allow to answer at least partially to the main questions posed in the introduction. First, the so-called *Niphargus stygius* group, following the definitions by Straškraba (1972), Vigna Taglianti (1972), and later by Karaman (1993) and Stoch (1998), is polyphyletic. Moreover, as already shown in a previous paper (Stoch et al. 2020), the group postulated by Vigna Taglianti (1972) for the Apennine populations (the so-called *speziae-romuleus-tatrensis* group) is polyphyletic as well, being the *N. tatrensis* clade of ancient origin and completely unrelated to the other two species which are part of the Apennine subclade of the main Italian clade. A polyphyly was already demonstrated by Delič et al. (2017) for the northwestern Balkan populations (Slovenia and NW Croatia) of the *N. stygius* group. The morphological similarities of all these different species (see Karaman, 1993 and Stoch, 1998, for a complete iconography for Italy), are maily based on: (i) their large body size (from 20 to 40 mm, the latter one reached by *N. costozzae*); (ii) large gnathopods and stout body shape; (iii) marked sexual dimorphism of the third uropod (elongated in males), and (iv) similar length of the two branches of the first male uropod (subequal as in female or only slightly different in length in males); (v) high similarity of mouthparts structure and setation and pereopods. These characters have proved to be inadequate for establishing phylogenetic relationships between different species, possibly due to convergence or retention of ancestral traits (Fišer et al. 2008). The incorrect interpretation of the value of these morphocharacters lead Karaman (1993) to attribute all the Italian populations belonging to the above-mentioned clades to a single species, i.e., *N. stygius*. Based on our results, it can be stated that the species *Niphargus stygius* is present in Italy only in the Dinaric Karst region of Trieste and Gorizia, and that a probable cryptic Alpine species (Delič et al. 2021) reaches the Julian Pre-Alps (Tab. S1, Uccea Valley), where *N. julius* occurs. The citations of *N. stygius* in caves on the Italian peninsula, especially in the ‘grey’ speleological literature, must therefore be considered unreliable.

Another species which should be probably deleted from the Italian fauna is *Niphargus forelii* (type locality: bottom of Lake Geneva, Switzerland) reported for Lake Tovel (Karaman and Ruffo 1993) and later for the Monte Baldo caves close to Garda Lake (Galassi et al., 2009). The specimens collected in these sites are included in the *Niphargus brixianus* clade, and only COI-based species delimitation methods consider them as a putative independent species. Finally, from a nomenclatorial point of view, the topotypes of *Niphargus brixianus* (Caja de Valmala, Brescia) are very similar to a population defined by Stoch (1998) as *Niphargus* cf. *lessiniensis* (Grotta Damati, Lessini Mountains, Verona), showing a faunistic affinity between the Prealps of Brescia and the Venetian Prealps, separated by the large Garda Lake and the Adige Valley. This paleogeographic scenario is confirmed by other taxa as well, like the specialized troglobitic genus *Lessinodytes* Vigna Taglianti, 1982 (Trechini ground beetles) present in both areas of which it is endemic (Faille et al., 2013). Another taxonomic problem is posed by the populations reported as *N. tridentinus* in GenBank (Bus Pursì, Brescia), part of the *N. brixianus* clades, and hence a misidentification (Sket, personal communication). The data downloaded from GenBank are incomplete; a short fragment (576 bp) of the COI gene suggests a close relationship with specimens of the *N. brixianus* clade present in the Bergamo Prealps, i.e., indicating a different scenario with a western affinity; even in this case there is an example in a troglobitic genus of Trechini beetles, *Allegrettia* Jeannel, 1928, present in both Bergamo and Brescia Prealps karstic areas of which it is endemic (Faille et al., 2013). Unfortunately, any effort by the authors to collect further material from that cave failed, due to flooding events in the area that probably obstructed its entrance; data are thus uncertain. It seems however possible that in the same area co-exist two haplogroups with different biogeographical affinities.

### Phylogeny and biogeography of niphargids in the Southern Limestone Alps

While the eastern clade (*Niphargus julius* clade) is quite problematic to allocate within the global phylogeny of the genus *Niphargus*, the other three alpine clades (*Niphargus brixianus, N. costozzae*, and *N. montellianus*) form a monophyletic group together with the Apennine *N. speziae* clade, suggestic they are Italian endemics. As regards the origin of the western Alpine clades, following the time-calibrated tree the split from the Apennine clade took place around 10 Ma (albeit the node support - posterior probability 0.79 - is not high enough to fully support this hypothesis), when the Apennine orogenesis came to an end and the possibility of colonization of the newly formed chain was realized. The low support of the node marking the separation between the *N. brixianus* and *N. speziae* clades does not allow to infer that the colonization of the Apennines started from the Alpine area.

A quick look at the distribution map reported in the present paper (Fig. 1) clearly shows that almost all populations of these clades inhabiting the Southern Limestone Alps are located south of the southern edge of the area occupied by glaciers during the last Last Glacial Maximum (LGM, around 21,000 ya), indicating that their evolutionary history is clearly pre-Pleistocenic, as strongly supported by the time-calibrated tree. The only exceptions are represented by the population of Lake Tovel (north of Garda Lake), and the one of *N. tridentinus*, settled in areas with an ice cover during the LGM. These populations can be considered as putative glacial relicts.

### Species richness and species delimitation methods

The results of the unilocus species delimitation methods applied in the present study (ASAP, PTP, and COI threshold) show a discordance in the number of putative species. Using the COI mitochondrial marker, estimates vary from 7 to 14 in the Western pre-Alpine clades and from 4 to 8 in the *N. julius* clade; looking at the time-calibrated tree, it seems plausible that separation of part of the putative species is due to the Pleistocenic glacial events. However, using the nuclear rDNA region, numbers are quite lower (4 putative species in the Western pre-Alpine clades and from 4 to 6 in the *N. julius* clade) showing a more conservative scenario.

Indeed, the mitonuclear discordance in SDMs results applied to nuclear and ribosomal markers is well known in niphargids (Stoch et al. 2020), especially in areas affected by glaciations. COI-based oversplitting may be quite common (Després 2019; Stoch et al., 2020) because of complex demographic history of lineage divergence/fusion due to the climatic fluctuations of the last million years. Relying on the fact that most of the recent splits in the Austrian clade of the *Niphargus tatrensis* complex took place during the Pleistocene, and different haplotypes were intermixed in the Northern Alps, Stoch et al. (2020) proposed to accept the species number suggested by the nuclear rDNA region. Albeit evidence of glaciations is widespread across Europe throughout the Quaternary (Ehlers and Gibbard, 2007), and the repeated glacial events may have isolated the different karstic massifs in different times, in the Italian Alps glaciation is identified possibly from the Plio/Pleistocene onwards but becomes established in the Southern Limestone Alps by MIS (Marine Isotope Stage) 22 only, i.e., 870–880 ka (Muttoni et al., 2012). These events are quite recent and their influence on speciation processes in niphargids virtually unknown: for this reason, to assign a specific status to the different populations clearly separated by glacial events is rather risky only on the basis of data from a single marker (COI).

Looking at both networks and trees, it seems evident that variability inside the *N. brixianus* clade in a small pre-Alpine area is higher than in the other clades, making difficult to take a definitive decision on the taxonomic status of its different populations. Some of these putative species seem to have been separated from the others in a pre-glacial period, suggesting that the western *N. brixianus* clade can be a complex of cryptic species. Pending further genetic analysis, we prefer to maintain a conservative approach here, and name all these populations as *Niphargus brixianus* s.l. (including the eastern morphospecies *N. lessiniensis* from the western Lessinian massif and *N*. cf. *forelii* from Monte Baldo massif and Tovel lake in Brenta dolomites). For the other clades, we suggest that most of the pre-glacial splits in the time-calibrated tree are indicative of the presence of different species, in agreement with nuclear rDNA-based species delimitation. For this reason, we confirm the subdivision of the *N. costozzae* clade in four species following the morphological results obtained by Stoch (1998), i.e., *N. costozzae* (widely distributed in the Venetian Pre-Alps), *N. tridentinus* (a putative glacial relict known only from few caves in southern Trentino and northern Venetian Pre-Alps), *N. montellianus* (Montello massif), and *N*. cf. *montellianus*, a nameless species distributed in eastern Venetian Pre-Alps and eastern Carnic Alps).

As regards the eastern clade, there is a sharp distinction between *N. julius* (Julian Pre-Alps), two new cryptic species (reported herein as *Niphargus* cf. *julius*) present respectively in the Carnic Pre-Alps and in the Carinthian Alps in Austria, and *N. kenki* from eastern Slovenian river basins (Sava River and its tributary, the Sotla River). The last species is the only one not found only in caves, but in alluvial sediments as well; however, it inhabits caves and springs (Karaman, 1952). Some species include distinct clades; the standard error of the common ancestors datation do not allow the exlusion of a separation due to glacial events from an older one. Furthermore, ‘hard’ splits may be a consequence of an insufficient sampling effort. Further research, following the example of the studies carried out on *Niphargus stygius* by Delić et al. (2021), will be able to shed light on the taxonomic significance of these cladogenetic events.

## Acknowledgements

Authors are indebted to the many speleologists who provided support and technical assistance in visiting complex caves, and to several speleologists and biologists who collected niphargid specimens during their excursions and made it available to us for study. We thank (in alphabetical order): Dante Bianco, Gianfranco Caoduro, Gianni Comotti, Luca Dorigo, Raoul Manenti, Erminio Piva, Gianfranco Tomasin, Vladimiro Toniello, and Dante Vailati.

## SUPPLEMENTARY MATERIAL

Figure S1 - Saturation plots mapping the observed pairwise divergences between indicidual COI sequences of Niphargidae against their genetic distance (p-value) for both transversions and transitions (saturation plots). Regression lines are fitted.

Figure S2 - Global maximum-likelihood phylogenetic tree of the genus *Niphargus* (using Pseudoniphargidae as outgroup) based on a concatenated dataset of ITC, 28S-I and COI (1st and 2nd codon only).

Figure S3 - ML phylogenetic trees based on nuclear rDNA (a, c) and mtDNA (b, d) markers of the two major *Niphargus* clades used for the PTP species delimitation method.

Table S1 - List of all the sequences included in the concatenated (ITS, 28S-I and COI) tree used in the analysis of the *Niphargus stygius* group. Sampling sites and WGS84 decimal degree coordinates are reported as well. (NEW = GebBank accession numbers will be added upon acceptance of the manuscript).

Table S2 - List of all the sequences downloaded from GenBank and integrated with new ones used, together with those listed in Tab. S1, to build the global tree of the family Niphargidae based on the concatenated dataset of nuclear (ITS+28S-I) and mitochondrial (COI, 1st+2nd codons only); three species of the sister family Pseudoniphargidae (*Microniphargus leruthi* and two species of the genus *Pseudoniphargus*) were used as an outgroup. (NEW = GebBank accession numbers will be added upon acceptance of the manuscript).

Table S3 - List of primers used for amplification and sequencing and PCR amplification conditions.

Table S4 - Results of the molecular species delimitation analysis using ASAP, PTP, and the patristic distance threshold methods.

