## Supplementary material for "Polyphyly of the *Niphargus stygius* species group (Crustacea, Amphipoda, Niphargidae) in the Southern Limestone Alps": Fig. S1

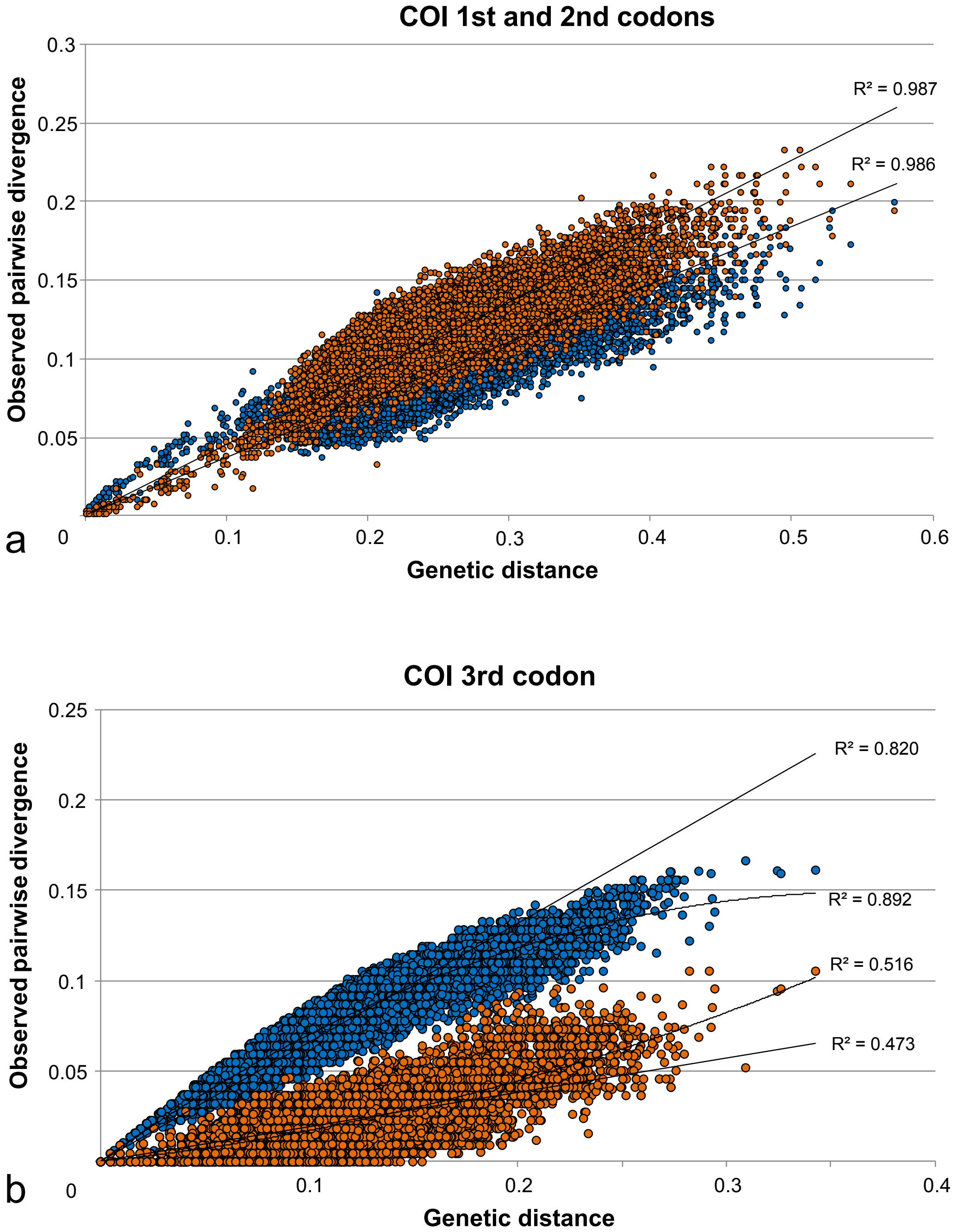


Figure S1 - Saturation plots mapping the observed pairwise divergences between indicidual COI sequences of Niphargidae against their genetic distance (p-value) for both transversions and transitions (saturation plots). Regression lines are fitted.
