## Supplementary material for "Polyphyly of the *Niphargus stygius* species group (Crustacea, Amphipoda, Niphargidae) in the Southern Limestone Alps": Fig. S2

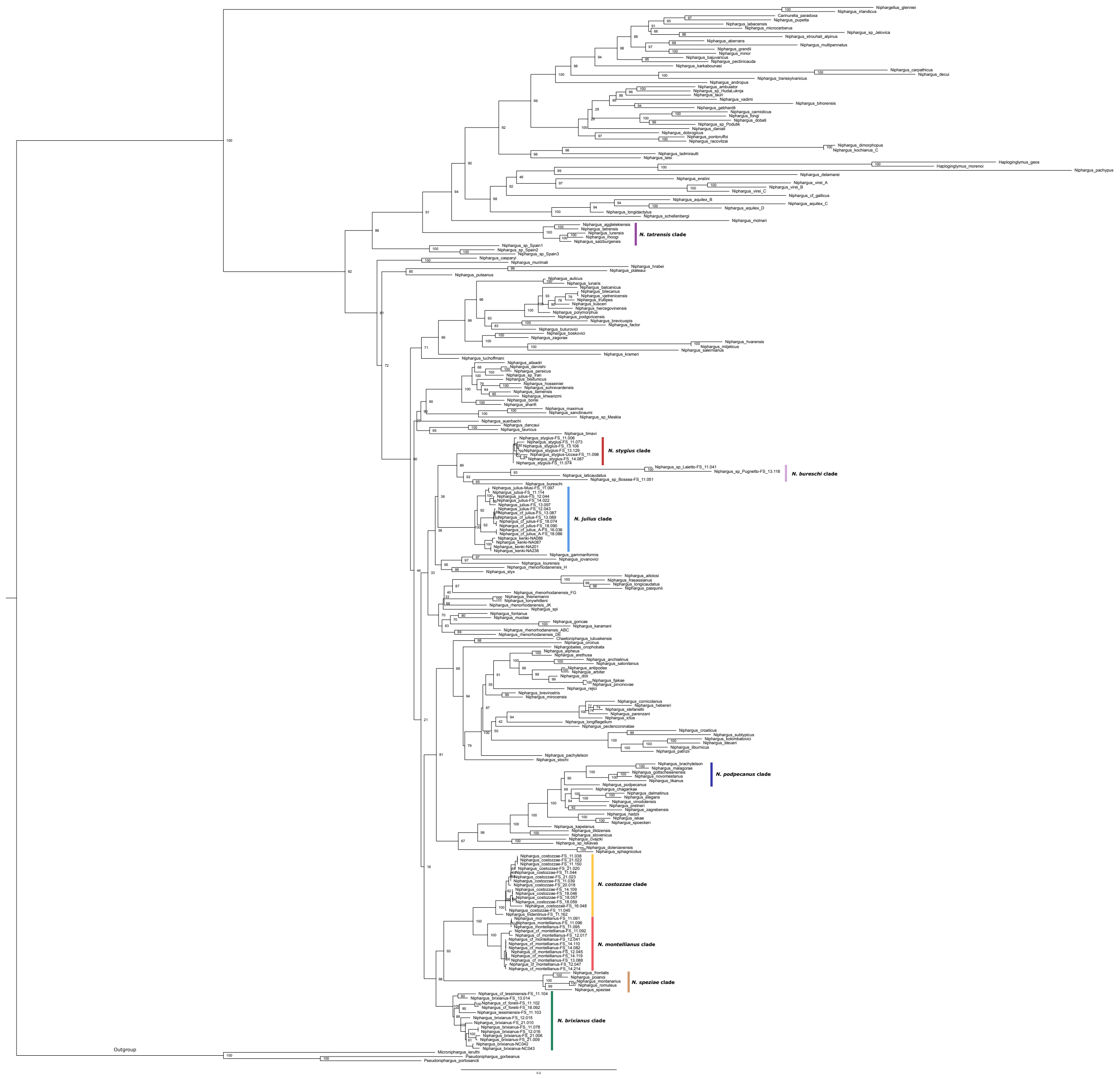

Figure S2 - Global maximum-likelihood phylogenetic tree of the genus *Niphargus* (using Pseudoniphargidae as outgroup) based on a concatenated dataset of ITC, 28S-I and COI (1st and 2nd codon only).
