## Supplementary material for "Polyphyly of the *Niphargus stygius* species group (Crustacea, Amphipoda, Niphargidae) in the Southern Limestone Alps": Fig. S3

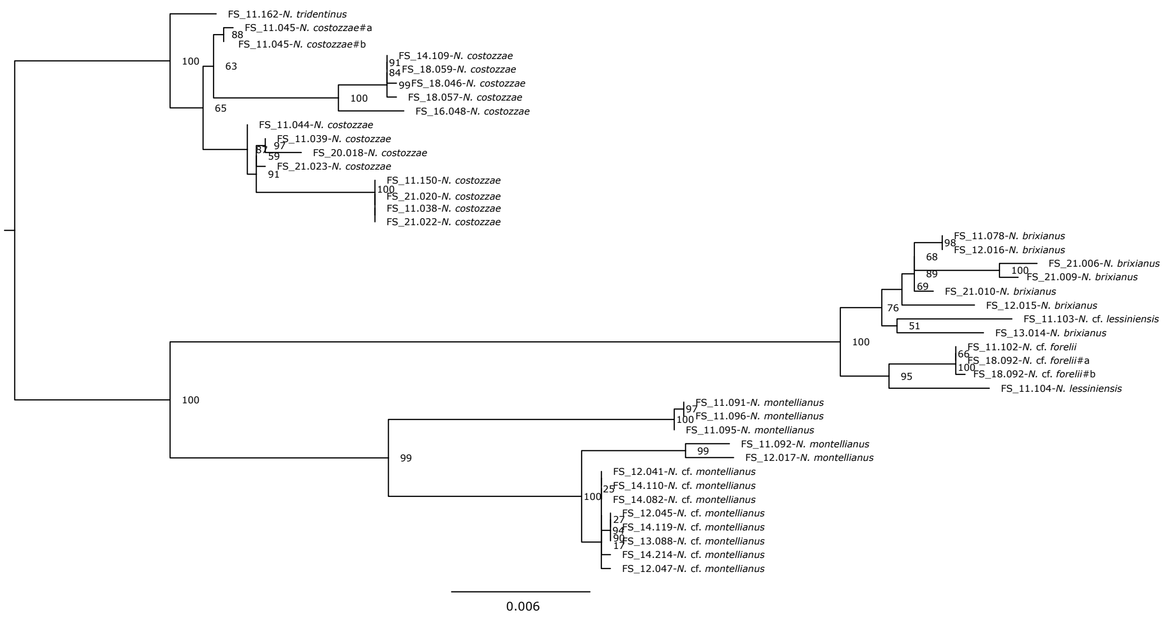


(a) ML phylogenetic tree of the western clade based on ITS+28S-I

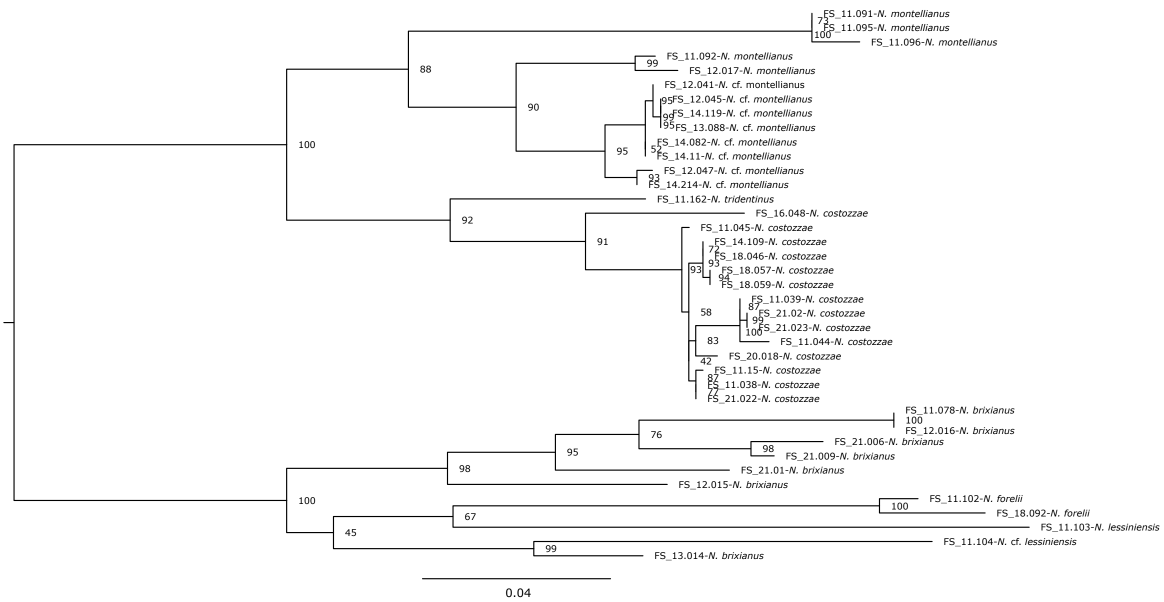


(b) ML phylogenetic tree of the western clade based on COI
