## Supplementary material for "Polyphyly of the *Niphargus stygius* species group (Crustacea, Amphipoda, Niphargidae) in the Southern Limestone Alps": Fig. S4

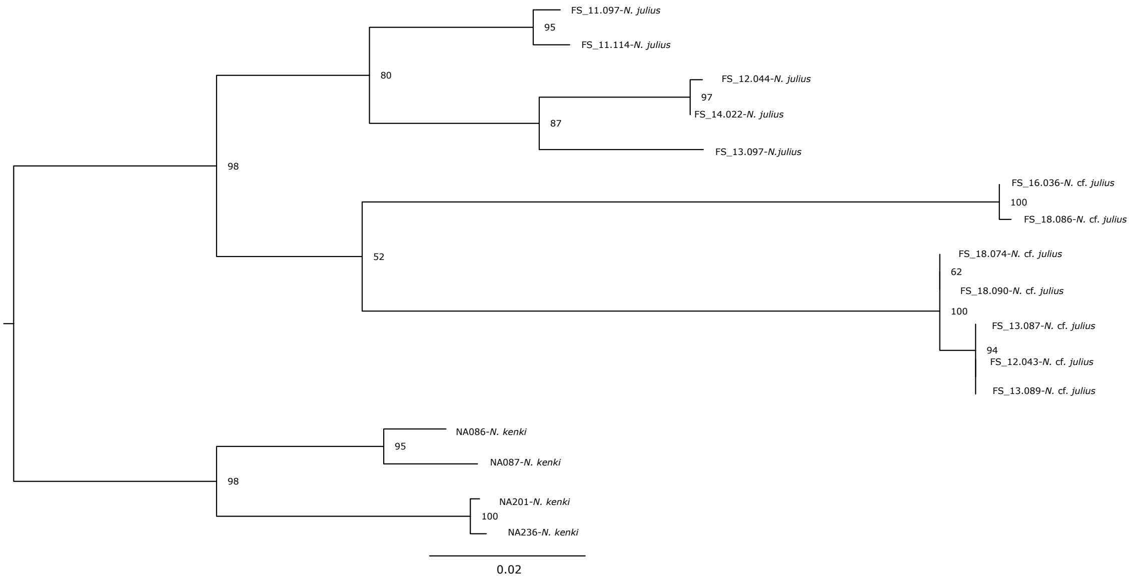


(c) ML phylogenetic tree of the eastern clade based on ITS+28S-I

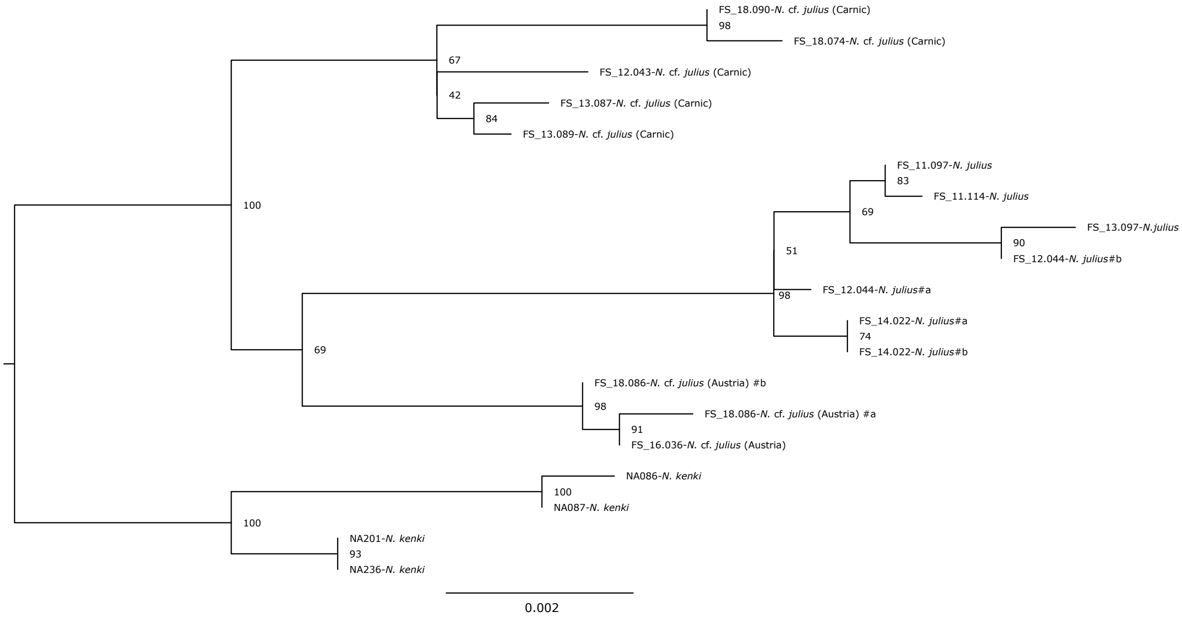


(d) ML phylogenetic tree of the eastern clade based on COI

Figure S3 - ML phylogenetic trees based on nuclear rDNA (a, c) and mtDNA (b, d) markers of the two major *Niphargus* clades used for the PTP species delimitation method.
