## Supplementary material for "Polyphyly of the *Niphargus stygius* species group (Crustacea, Amphipoda, Niphargidae) in the Southern Limestone Alps": Tab. S1

Table S1 - List of all the sequences included in the concatenated (ITS, 28S-I and COI) tree used in the analysis of the *Niphargus stygius* species group. Sampling sites and WGS84 decimal degree coordinates are reported as well. (NEW = GebBank accession numbers will be added upon acceptance of the manuscript).

| Voucher | Species | Country | Region | Site | Longitude | Latitude | ITS | 28S | COI |
|---|---|---|---|---|---|---|---|---|---|
| FS_11.006 | *Niphargus stygius* | Italy | Friuli Venezia Giulia | Grotta Azzurra di Samatorza (VG 257) | 13.705222 | 45.752871 | NEW | MT975470 | KY706707 |
| FS_11.038 | *Niphargus costozzae* | Italy | Veneto | Buso del Becco d'Oro (154 V/VI), S. Tomio (Monti Lessini) | 11.420376 | 45.64229 | NEW | NEW | KY706712 |
| FS_11.039 | *Niphargus costozzae* | Italy | Veneto | Buso delle Anguane (518 V/VI) (Monti Lessini) | 11.277764 | 45.626234 | NEW | NEW | KY706713 |
| FS_11.044 | *Niphargus costozzae* | Italy | Veneto | Grotta della Guerra (127 V/VI), Lumignano (Monti Berici) | 11.578398 | 45.455608 | NEW | NEW | KY706718 |
| FS_11.045a | *Niphargus costozzae* | Italy | Veneto | Cogolo delle Tette (36 V/VI), Val Pizzarda (Monti Berici) | 11.42746 | 45.384581 | NEW | NEW | KY706719 |
| FS_11.045b | *Niphargus costozzae* | Italy | Veneto | Cogolo delle Tette (36 V/VI), Val Pizzarda (Monti Berici) | 11.42746 | 45.384581 | NEW | NEW | KY706719 |
| FS_11.073 | *Niphargus stygius* | Italy | Friuli Venezia Giulia | Galleria di guerra tra Devetachi e Visintini | 13.568209 | 45.867281 | NEW | NEW | KY706880 |
| FS_11.074 | *Niphargus stygius* | Italy | Friuli Venezia Giulia | Sorgente Bukovec superiore, Val Rosandra | 13.872203 | 45.614048 | NEW | NEW | KY706881 |
| FS_11.078 | *Niphargus brixianus* | Italy | Lombardia | Buco del Corno o Büs del Córen (1247 Lo/BG) | 9.534333 | 45.78982 | NEW | NEW | KY706885 |
| FS_11.091 | *Niphargus montellianus* | Italy | Veneto | Tavaran Grande (69 V/TV), Campagnole di Sopra (Montello) | 12.148766 | 45.845068 | NEW | KT878856 | KY706887 |
| FS_11.092 | *Niphargus cf. montellianus* | Italy | Veneto | Busa delle Fave (1272 V/TV), Cà Brisotto (Colli di Conegliano) | 12.213267 | 45.928052 | NEW | NEW | KY706888 |
| FS_11.095 | *Niphargus montellianus* | Italy | Veneto | Grotta Crede-a (2098 V/TV) (Colli di Conegliano) | 12.206488 | 45.879319 | NEW | NEW | KY706721 |
| FS_11.096 | *Niphargus montellianus* | Italy | Veneto | Buoro di Ciano (71 V/TV), Santa Margherita (Montello) | 12.087052 | 45.633947 | NEW | NEW | KY706722 |
| FS_11.097 | *Niphargus julius* | Italy | Friuli Venezia Giulia | Sorgente sul sentiero 727 (Fr 2786), Monte Musi | 13.330599 | 46.330528 | NEW | NEW | KY706723 |
| FS_11.098 | *Niphargus stygius* | Italy | Friuli Venezia Giulia | Sorgente del Torrente Uccea | 13.328411 | 46.334873 | NEW | NEW | KY706724 |
| FS_11.102 | *Niphargus* cf. *forelii* | Italy | Veneto | Grotta dei Trovai (157 V/VR), Casara Trovai (Monte Baldo) | 10.796595 | 45.696289 | NEW | NEW | KY706728 |
| FS_11.103 | *Niphargus lessiniensis* | Italy | Veneto | Grotta A del Ponte di Veja (117 V/VR), Veja (Monti Lessini) | 10.970374 | 45.608447 | NEW | NEW | KY706729 |
| FS_11.104 | *Niphargus* cf. *lessiniensis* | Italy | Veneto | Grotta dei Damati (9 V/VR), Case Damati (Monti Lessini) | 11.163264 | 45.560788 | NEW | NEW | KY706730 |
| FS_11.114 | *Niphargus julius* | Italy | Friuli Venezia Giulia | Grotta risorgiva di Starcedat (Fr 483) | 13.53733 | 46.108667 | NEW | NEW | KY706739 |
| FS_11.150 | *Niphargus costozzae* | Italy | Veneto | Grotta della Poscola (136 V/VI), Priabona (Monti Lessini) | 11.370682 | 45.633317 | NEW | NEW | KY706752 |
| FS_11.162 | *Niphargus tridentinus* | Italy | Trentino | Grotta della Bigonda (243 VT/TN), Selva di Grigno | 11.580878 | 46.018206 | NEW | KT878857 | KY706896 |
| FS_12.015 | *Niphargus brixianus* | Italy | Lombardia | Cava (miniera) di Pietre Coti, località Pratberta | 9.748595 | 45.747447 | NEW | NEW | KY706899 |
| FS_12.016 | *Niphargus brixianus* | Italy | Lombardia | Grotta dè Val D'Adda (1044 Lo/BG), Ca Contaglio | 9.525603 | 45.801263 | NEW | NEW | KY706900 |
| FS_12.017 | *Niphargus* cf. *montellianus* | Italy | Veneto | Grotte del Caglieron, sorgente nelle cave di arenaria | 12.330928 | 46.007097 | NEW | NEW | KY706901 |
| FS_12.041 | *Niphargus* cf. *montellianus* | Italy | Friuli Venezia Giulia | Risorgiva 1a a W di Ominutz (2760 Fr) | 12.884729 | 46.241661 | NEW | NEW | KY706912 |
| FS_12.043 | *Niphargus* cf. *julius* | Italy | Friuli Venezia Giulia | Caverna Mainarda o Mainarie dal Puint (Fr 242), Pradis | 12.88399 | 46.247179 | NEW | NEW | KY706913 |
| FS_12.044a | *Niphargus julius* | Italy | Friuli Venezia Giulia | Grotta Doviza (Fr 70), Villanova | 13.287064 | 46.251771 | NEW | NEW | KY706914 |
| FS_12.044b | *Niphargus julius* | Italy | Friuli Venezia Giulia | Grotta Doviza (Fr 70), Villanova | 13.287064 | 46.251771 | NEW | NEW | KY706914 |
| FS_12.045 | *Niphargus* cf. *montellianus* | Italy | Friuli Venezia Giulia | Sorgenti Pissui, valle del Torrente Colvera | 12.70572 | 46.18424 | NEW | NEW | KY706915 |
| FS_12.047 | *Niphargus* cf. *montellianus* | Italy | Friuli Venezia Giulia | Sorgenti del Fiume Livenza La Santissima | 12.475922 | 46.021297 | NEW | NEW | KY706917 |
| FS_13.014 | *Niphargus brixianus* | Italy | Lombardia | Caìa de Valmala (141 Lo/BS), Marcheno | 10.234563 | 45.716559 | NEW | NEW | NEW(3) |
| FS_13.087 | *Niphargus* cf. *julius* | Italy | Friuli Venezia Giulia | Caverna Mainarda o Mainarie dal Puint (Fr 242), Pradis | 12.88399 | 46.247179 | NEW | NEW | KY706772 |
| FS_13.088 | *Niphargus* cf. *montellianus* | Italy | Friuli Venezia Giulia | Fornat di Meduno (Fr 123) | 12.776542 | 46.222099 | NEW | NEW | KY706773 |
| FS_13.089 | *Niphargus* cf. *julius* | Italy | Friuli Venezia Giulia | Sorgente presso Pielungo | 12.926671 | 46.273914 | NEW | NEW | KY706774 |
| FS_13.097 | *Niphargus julius* | Italy | Friuli Venezia Giulia | Grotta di Papipano (Fr 296), Canal di Grivò | 13.357085 | 46.162098 | NEW | NEW | KY706780 |
| FS_13.106 | *Niphargus stygius* | Italy | Friuli Venezia Giulia | Sorgenti di Moschenizze | 13.584766 | 45.801616 | NEW | NEW | KY706788 |
| FS_13.129 | *Niphargus stygius* | Italy | Friuli Venezia Giulia | Sorgente presso ramo I del Timavo | 13.591139 | 45.787724 | NEW | NEW | KY706809 |
| FS_14.022a | *Niphargus julius* | Italy | Friuli Venezia Giulia | Grotta Nuova di Villanova (Fr 323), Villanova Grotte | 13.282433 | 46.257155 | NEW | NEW | KY706943 |
| FS_14.022b | *Niphargus julius* | Italy | Friuli Venezia Giulia | Grotta Nuova di Villanova (Fr 323), Villanova Grotte | 13.282433 | 46.257155 | NEW | NEW | KY706943 |
| FS_14.082 | *Niphargus* cf. *montellianus* | Italy | Friuli Venezia Giulia | Grotta della Foos (Fr 229), Campone | 12.813839 | 46.254493 | NEW | NEW | KY706960 |
| FS_14.087 | *Niphargus stygius* | Italy | Friuli Venezia Giulia | Sorgente Skedanz, Santa Croce | 13.696698 | 45.724674 | NEW | NEW | KY706964 |
| FS_14.109 | *Niphargus costozzae* | Italy | Veneto | Fonte nella piazza di Fontanafredda, Colli Euganei | 11.661603 | 45.291217 | NEW | NEW | NA |
| FS_14.110 | *Niphargus* cf. *montellianus* | Italy | Friuli Venezia Giulia | Grotta della Foos (Fr 229), Campone | 12.813839 | 46.254493 | NEW | NEW | KY706978 |
| FS_14.119 | *Niphargus* cf. *montellianus* | Italy | Friuli Venezia Giulia | Sorgente a Rovareit | 12.73692 | 46.199446 | NEW | NEW | KY706987 |
| FS_14.214 | *Niphargus* cf. *montellianus* | Italy | Friuli Venezia Giulia | Sorgente vecchia strada Valcellina | 12.603927 | 46.186851 | NEW | NEW | KY706825 |
| FS_16.036 | *Niphargus* cf. *julius* | Austria | Kärnten | Obir Tropfsteinhöhle (3925/3), Bad Eisenkappel, Kärnten | 14.545 | 46.509167 | NEW | NEW | NEW |
| FS_16.048 | *Niphargus costozzae* | Italy | Veneto | Grotta del Nespolo del Mare (3140 V/VI), Vesene | 11.531794 | 45.769566 | NEW | NEW | NA |
| FS_18.046 | *Niphargus costozzae* | Italy | Veneto | Pozzo a Fontanafredda | 11.670516 | 45.291117 | NEW | NEW | NEW |
| FS_18.057 | *Niphargus costozzae* | Italy | Veneto | Buso della Casara, Valnogaredo, Colli Euganei | 11.67323 | 45.30747 | NEW | NEW | NEW |
| FS_18.059 | *Niphargus costozzae* | Italy | Veneto | Fontana nella piazza di Fontanafredda, Colli Euganei | 11.661206 | 45.29126 | NEW | NEW | NEW |
| FS_18.074 | *Niphargus* cf. *julius* | Italy | Friuli Venezia Giulia | Pitfall trap Col Palotta, Solimbergo | 12.833092 | 46.184349 | NEW | NEW | NEW |
| FS_18.086a | *Niphargus* cf. *julius* | Austria | Kärnten | Small spring in gorge Kupitzlamm, Bad Eisenkappel | 14.615833 | 46.464722 | NEW | NEW | NEW |
| FS_18.086b | *Niphargus* cf. *julius* | Austria | Kärnten | Small spring in gorge Kupitzlamm, Bad Eisenkappel | 14.615833 | 46.464722 | NEW | NEW | NEW |
| FS_18.090 | *Niphargus* cf. *julius* | Italy | Friuli Venezia Giulia | Grotta presso Solimbergo (3519 Fr) | 12.8373 | 46.1858 | NEW | NEW | NEW |
| FS_18.092 | *Niphargus* cf. *forelii* | Italy | Trentino | Sorgenti del Torrente Tresenga, Valle di Tovel | 10.975734 | 46.29286 | NEW | NEW | NEW |
| FS_20.018 | *Niphargus costozzae* | Italy | Veneto | Grottina-Sorgente Valle Anguane, Recoaretto, Castelcerino | 11.232844 | 45.457547 | NEW | NEW | NEW |
| FS_21.009 | *Niphargus* cf. *brixianus* | Italy | Lombardia | Miniera Pelucchi | 9.403406 | 45.732591 | NEW | NEW | NEW |
| FS_21.010 | *Niphargus* cf. *brixianus* | Italy | Lombardia | Risorgente dei Camosci | 9.493889 | 45.879925 | NEW | NEW | NEW |
| FS_21.020 | *Niphargus costozzae* | Italy | Veneto | Fontana Zordan, Parigi | 11.413087 | 45.62049 | NEW | NEW | NEW |
| FS_21.022 | *Niphargus costozzae* | Italy | Veneto | Sorgente Pedrina, Torreselle | 11.425011 | 45.605447 | NEW | NEW | NEW |
| FS_21.023 | *Niphargus costozzae* | Italy | Veneto | Fontana de Zaicamomo, Ignago | 11.44933 | 45.605447 | NEW | NEW | NEW |
| NA086 | *Niphargus kenki* | Slovenia |  | Izvir pod hišo Sodna vas 25, Sodna vas, Podčetrtek | 15.594861 | 46.176186 | KY617662 | KR905869 | KR905804 |
| NA087 | *Niphargus kenki* | Slovenia |  | Polje of Sotli, Bistrica ob Sotli, Brežice | 15.6641 | 46.0679 | KY617663 | EU693306 | KY617476 |
| NA201 | *Niphargus kenki* | Slovenia |  | Marijino brezno, Lubnik, Škofja loka | 14.296336 | 46.164652 | KY617669 | KY617368 | KY617479 |
| NA236 | *Niphargus kenki* | Slovenia |  | Ljubljanica river, phreatic water, Tomačevo, Ljubljana | 14.54057 | 46.08281 | KY617670 | KY617369 | KY617480 |
