## Supplementary material for "Polyphyly of the *Niphargus stygius* species group (Crustacea, Amphipoda, Niphargidae) in the Southern Limestone Alps": Tab. S2

Table S2 - List of all the sequences downloaded from GenBank and integrated with new ones used, together with those listed in Tab. S1, to build the global tree of the family Niphargidae based on the concatenated dataset of nuclear (ITS+28S-I) and mitochondrial (COI, 1st+2nd codons only) markers; three species of the sister family Pseudoniphargidae (*Microniphargus leruthi* and two species of the genus *Pseudoniphargus*) were used as an outgroup. (NEW = GebBank accession numbers will be added upon acceptance of the manuscript).

| **Species** | **Voucher** | **ITS** | **28S** | **COI** |
|---|---|---|---|---|
| *Carinurella paradoxa* | NA738 | NA | OK030552 | KR905829 |
| *Chaetoniphargus lubuskensis* | ND831 | NA | OK157634 | OK157253 |
| *Haploginglymus geos* | PN1374 | NA | KY441086 | NA |
| *Haploginglymus morenoi* | PN0243 | NA | KY441079 | KY441020 |
| *Microniphargus leruthi* | DW190413-008 | NA | MT994441 | MT993552 |
| *Niphargellus glenniei* | NC017 | NA | MN914041 | NA |
| *Niphargobates orophobata* | NA547 | NA | JQ815552 | KY643582 |
| *Niphargus aberrans* | ND600 | **NEW** | OK157525 | OK157092 |
| *Niphargus aggtelekiensis* | FS_14.016 | **NEW** | MT975473 | KY706939 |
| *Niphargus aitolosi* | NA081 | NA | EU693310 | MN113979 |
| *Niphargus alisadri* | NB105 | NA | KF581049 | KF581079 |
| *Niphargus alpheus* | NB169 | KY617323 | KY617132 | KY617251 |
| *Niphargus ambulator* | NA504 | NA | KJ566699 | KX379125 |
| *Niphargus anchialinus* | NB168 | KY617349 | KR905881 | KY617287 |
| *Niphargus andropus* | NA942 | NA | KF218725 | NA |
| *Niphargus antipodes* | NB604 | KY617318 | KY617131 | KY643566 |
| *Niphargus aquilex* B | NA020 | NA | EF617255 | KX379140 |
| *Niphargus aquilex* C | GBLP1 | NA | JF420876 | JF420842 |
| *Niphargus aquilex* D | UK | NA | KC315607 | KC315622 |
| *Niphargus arbiter* | NB419 | KY617328 | KY617099 | KY617260 |
| *Niphargus arethusa* | NA051 | KY617332 | EF617285 | KY617264 |
| *Niphargus auerbachi* | NA069 | NA | EU693292 | KX379130 |
| *Niphargus aulicus* | NC099 | NA | MN914026 | MN913980 |
| *Niphargus bajuvaricus* | NA024 | NA | EF617259 | KX379135 |
| *Niphargus balcanicus* | NA070 | NA | EF617280 | MN913981 |
| *Niphargus bihorensis* | NA792 | NA | KF218727 | KF218663 |
| *Niphargus bilecanus* | NA182 | NA | JQ815550 | KR905826 |
| *Niphargus bisitunicus* | NB113 | NA | KF581050 | KF581062 |
| *Niphargus borisi* | NB123 | NA | KF581044 | KF581076 |
| *Niphargus boskovici* | NA036 | NA | EF617271 | KR905781 |
| *Niphargus brachytelson* | NA071 | NA | EU693293 | KR905797 |
| *Niphargus brevicuspis* | NC034 | NA | MN914028 | KY643550 |
| *Niphargus brevirostris* | NC123 | NA | MN914008 | NA |
| *Niphargus bureschi* | NA164 | NA | MN114020 | KY643614 |
| *Niphargus buturovici* | NC032 | NA | MN914022 | KY643551 |
| *Niphargus carniolicus* | NA017 | NA | EF617252 | KR905776 |
| *Niphargus carpathicus* | NA149 | NA | MN114019 | KY643616 |
| *Niphargus casparyi* | NA525 | NA | KX379003 | KX379123 |
| *Niphargus* cf *gallicus* | NA145 | NA | MN914040 | KY643633 |
| *Niphargus chagankae* | NB794 | KY617689 | KY617399 | KY617499 |
| *Niphargus cornicolanus* | FS_14.029 | NA | MN914003 | MN913987 |
| *Niphargus croaticus* | NB013 | KT007275 | KT007482 | KT007342 |
| *Niphargus cvajcki* | NB800 | KY617695 | KY617405 | KY617505 |
| *Niphargus dalmatinus* | NA060 | KY617657 | EF617296 | KR905790 |
| *Niphargus dancaui* | CF_10.4 | KF290133 | MN914033 | MN913990 |
| *Niphargus daniali* | NB132 | NA | KF581033 | KF581080 |
| *Niphargus darvishi* | NB117 | NA | KF581043 | KF581072 |
| *Niphargus decui* | NA154 | KF290197 | KF719272 | KR905822 |
| *Niphargus delamarei* | NA075 | NA | EU693295 | NA |
| *Niphargus dimorphopus* | NA136 | NA | KJ566702 | KX379138 |
| *Niphargus dobati* | NA013 | NA | EF617247 | KR905774 |
| *Niphargus dobrogicus* | NA140 | NA | KR905871 | KR905816 |
| *Niphargus dolenianensis* | NA034 | NA | EF617269 | KX379133 |
| *Niphargus doli* | NB173 | KY617324 | KR905882 | KY617255 |
| *Niphargus elegans* | NA061 | KY617658 | EF617297 | KR905791 |
| *Niphargus enslini* | DW170827-01 | NA | OK030554 | OK021662 |
| *Niphargus factor* | NA078 | NA | EU693298 | KR905798 |
| *Niphargus fjakae* | NA054 | KY617315 | EF617290 | KY617285 |
| *Niphargus fongi* | NA538 | NA | MN914037 | KY643584 |
| *Niphargus fontanus* | NA526 | **NEW** | KJ566686 | KX379122 |
| *Niphargus frasassianus* | NC027 | GU973409 | GU973414 | GU973144 |

| | | | | |
|---|---|---|---|---|
| *Niphargus frontalis* (clade 4) | LPR_08.3 | GU973272 | GU973423 | GU973053 |
| *Niphargus gammariformis* | ME_13.56 | NA | MN114013 | KY707025 |
| *Niphargus gebhardti* | NB550 | NA | KP967556 | KP967553 |
| *Niphargus goricae* | NB932 | KY617724 | KY617447 | KY617534 |
| *Niphargus gottscheeanensis* | NB844 | KY617699 | KY617412 | KY617512 |
| *Niphargus grandii* | NA080 | NA | EU693300 | KR905799 |
| *Niphargus hadzii* | NA082 | KY617660 | EU693301 | KR905800 |
| *Niphargus hebereri* | BR_10.1 | NA | MT133569 | KY706634 |
| *Niphargus hercegovinensis* | NA151 | NA | JQ815549 | KR905820 |
| *Niphargus hosseiniei* | NB106 | NA | KF581054 | KF581064 |
| *Niphargus hrabei* | NB535 | KF290147 | EU693302 | MG548163 |
| *Niphargus hvarensis* | NA038 | NA | JQ815452 | KR905782 |
| *Niphargus ictus* | NC026 | GU973323 | KX379008 | KX379110 |
| *Niphargus ilamensis* | NB111 | NA | KF581038 | KF581069 |
| *Niphargus illidzensis* | NA084 | KY617661 | EU693304 | KR905802 |
| *Niphargus irlandicus* | NC013 | NA | KC315618 | KY643557 |
| *Niphargus iskae* | NB619 | KY617731 | KY617382 | KY617490 |
| *Niphargus jovanovici* | ME_13.100 | NA | MN114014 | KY707023 |
| *Niphargus kapelanus* | NB625 | NA | KY617387 | KY617494 |
| *Niphargus karamani* | NB933 | NA | EU693305 | KR905803 |
| *Niphargus karkabounasi* | NA217 | NA | KR905877 | KP164477 |
| *Niphargus khwarizmi* | NB119 | NA | KF581057 | KF581063 |
| *Niphargus kochianus* C | UK_HT13 | NA | KC315612 | KC315675 |
| *Niphargus kolombatovici* | NC826 | KT007241 | OK157545 | OK156624 |
| *Niphargus krameri* | FS_13.098 | **NEW** | **NEW** | KY706781 |
| *Niphargus kusceri* | NB422 | NA | KR905883 | MZ270543 |
| *Niphargus labacensis* | NA022 | NA | EF617257 | KR905777 |
| *Niphargus ladmiraulti* | DM_lad1 | NA | GU973462 | GU973424 |
| *Niphargus laisi* | NA135 | NA | KX379002 | KX379076 |
| *Niphargus laticaudatus* | NA905 | NA | KF218722 | KF218687 |
| *Niphargus lessiniensis* | NA064 | NA | EF617300 | KR905794 |
| *Niphargus liburnicus* | NB134 | **NEW** | KT007477 | KT007419 |
| *Niphargus likanus* | NB917 | KY617713 | KY617435 | KY617531 |
| *Niphargus longicaudatus* | CF_09.05 | **NEW** | **NEW** | KY706646 |
| *Niphargus longidactylus* | NA541 | NA | EF617256 | KY643583 |
| *Niphargus longiflagellum* | NB205 | KR905840 | MN914006 | NA |
| *Niphargus lourensis* | NA094 | NA | EU693312 | KR905806 |
| *Niphargus luchoffmani* | NC204 | MH172382 | MH172405 | MH172433 |
| *Niphargus lunaris* | NC792 | NA | OK157641 | OK156591 |
| *Niphargus lurensis* | FS_18.077 | MT975505 | MT975505 | MT975663 |
| *Niphargus malagorae* | NB621 | KY617682 | KY617384 | KY643562 |
| *Niphargus maximus* | NA044 | NA | EF617279 | KY643603 |
| *Niphargus microcerberus* | NA600 | NA | MN114023 | KY643575 |
| *Niphargus miljeticus* | NA500 | NA | KR905878 | NA |
| *Niphargus minor* | NC031 | NA | MN114028 | KY643552 |
| *Niphargus mirocensis* | NB909 | NA | KR827047 | NA |
| *Niphargus molnari* | NB554 | NA | MN914038 | KY643567 |
| *Niphargus montanarius* | NC025 | GU973222 | GU973421 | GU973018 |
| *Niphargus moogi* | FS_17.013 | MT975462 | MT975499 | MT975657 |
| *Niphargus multipennatus* | NA169 | NA | KJ566700 | KR905825 |
| *Niphargus muotae* | NC061 | NA | KX379024 | KX379095 |
| *Niphargus murimali* | NC059 | NA | KX379022 | KX379097 |
| *Niphargus novomestanus* | NA096 | KY617740 | EU693314 | KY617552 |
| *Niphargus orcinus* | NA099 | KR905842 | EU693315 | KR905808 |
| *Niphargus pachypus* | FS_14.158 | NA | **NEW** | KY706994 |
| *Niphargus pachytelson* | NB420 | KR905841 | EU693316 | KR905809 |
| *Niphargus parenzani* | FS_11.120 | NA | MN913997 | KY706745 |
| *Niphargus pasquinii* | CF_09.06 | **NEW** | EF617244 | MN913989 |
| *Niphargus patrizii* | FS_13.016 | NA | MN914011 | KY706926 |
| *Niphargus pectencoronatae* | FS_14.229 | NA | MN914010 | KY706831 |
| *Niphargus pectinicauda* | NA023 | NA | EF617258 | KR905778 |
| *Niphargus persicus* | NB128 | NA | KF581047 | KF581075 |
| *Niphargus pincinovae* | NB471 | KY617351 | KR905884 | KY617283 |
| *Niphargus plateaui* | DM_pl1 | NA | EF025851 | GU973427 |
| *Niphargus podgoricensis* | NA166 | NA | KR905875 | KR905824 |
| *Niphargus podpecanus* | NB910 | KY617709 | KY617430 | KY617526 |
| *Niphargus poianoi* | NC024 | **NEW** | KX379006 | KX379112 |
| *Niphargus polymorphus* | NA047 | NA | EF617282 | KR905784 |
| *Niphargus pontoruffoi* | NB093 | NA | MK421144 | KY643637 |
| *Niphargus pretneri* | NC101 | NA | MT191525 | MT192023 |
| *Niphargus pupetta* | FS_14.026 | NA | **NEW** | KY706946 |
| *Niphargus puteanus* | NA066 | NC162 | EF617302 | KR905795 |

| | | | | |
|---|---|---|---|---|
| *Niphargus racovitzai* | NB094 | NA | MN914036 | KY707072 |
| *Niphargus rejici* | NA048 | KR905833 | EF617283 | KR905785 |
| *Niphargus rhenorhodanensis* ABC | NB437 | NA | KJ566681 | KX379117 |
| *Niphargus rhenorhodanensis* DE | NA104 | NA | EU693319 | KR905811 |
| *Niphargus rhenorhodanensis* FG | NC046 | NA | KX379034 | KX379107 |
| *Niphargus rhenorhodanensis* H | NB439 | NA | KJ566685 | KX379116 |
| *Niphargus rhenorhodanensis* JK | NC219 | NA | MH172416 | MH172436 |
| *Niphargus romuleus* | NC033 | **NEW** | MT975475 | KY706950 |
| *Niphargus salernianus* | FS_10.011 | NA | MN914014 | KY706696 |
| *Niphargus salonitanus* | NA053 | KR905839 | EF617289 | KR905788 |
| *Niphargus salzburgensis* | FS_14.179 | MT975453 | MT975480 | KY707001 |
| *Niphargus sanctinaumi* | NA105 | NA | EU693320 | KR905812 |
| *Niphargus schellenbergi* | NA032 | GU973481 | EF617267 | KR905780 |
| *Niphargus sharifi* | NB118 | NA | KF581048 | KF581078 |
| *Niphargus slovenicus* | ND149 | KY617666 | OK157565 | OK156731 |
| *Niphargus sohrevardensis* | NB112 | NA | KF581034 | KF581061 |
| *Niphargus* sp. HudaLuknja | NA012 | NA | EF617246 | KY643627 |
| *Niphargus* sp. Iran | NB125 | NA | KF581040 | KR905831 |
| *Niphargus* sp. Iskavas | NA055 | NA | EF617291 | KY643623 |
| *Niphargus* sp. Jelovica | NA515 | NA | MN114022 | KY643586 |
| *Niphargus* sp. Meskla | NC213 | NA | MN114031 | MN113981 |
| *Niphargus* sp. Podutik | NA016 | NA | EF617251 | KY643626 |
| *Niphargus* sp. Spain1 | PN1259 | NA | KY441088 | KY441035 |
| *Niphargus* sp. Spain2 | PN0250 | NA | KY441083 | KY441030 |
| *Niphargus* sp. Spain3 | PN0521 | NA | KY441096 | KY441025 |
| *Niphargus speziae* | FS_11.069 | **NEW** | MT975471 | KY706876 |
| *Niphargus sphagnicolus* | NA035 | NA | EF617270 | KR858495 |
| *Niphargus spinulifemur* | NB187 | **NEW** | **NEW** | KY706878 |
| *Niphargus spoeckeri* | NA108 | KY617667 | EU693324 | KR905814 |
| *Niphargus stefanellii* | FS_11.072 | **NEW** | MN913999 | KY706879 |
| *Niphargus steueri* | NB041 | KT007229 | KT007496 | KT007368 |
| *Niphargus stochi* | NA599 | NA | JQ815551 | KY643576 |
| *Niphargus strouhali alpinus* | FS_11119 | NA | EF617254 | KY706744 |
| *Niphargus styx* | NC060 | NA | KX379023 | KX379096 |
| *Niphargus subtypicus* | NA112 | KT007230 | EU693326 | KT007433 |
| *Niphargus tatrensis* | NC106 | **NEW** | MT975496 | MT975654 |
| *Niphargus tauri* | NA011 | NA | EF617245 | NA |
| *Niphargus tauricus* | NA155 | NA | KR905874 | KR905823 |
| *Niphargus thienemanni* | NB443 | MH172396 | KJ566688 | KX379114 |
| *Niphargus timavi* | NA190 | **NEW** | MN914034 | MN913985 |
| *Niphargus tonywhitteni* | NC178 | MH172398 | MH172414 | MH172432 |
| *Niphargus transsylvanicus* | NA904 | NA | KF218733 | KF218715 |
| *Niphargus trullipes* | NA046 | NA | EF617281 | KR905783 |
| *Niphargus vadimi* | NA144 | NA | KR905872 | KR905817 |
| *Niphargus vinodolensis* | NA062 | KY617659 | EF617298 | KR905792 |
| *Niphargus virei* A | FD9 | **NEW** | DQ119313 | DQ064715 |
| *Niphargus virei* B | NB402 | NA | KJ566680 | KX379098 |
| *Niphargus virei* C | NA003 | NA | EF617237 | KR905771 |
| *Niphargus vjetrenicensis* | NA116 | KR905844 | EU693329 | MT192020 |
| *Niphargus zagorae* | NA212 | NA | KR827044 | KR827046 |
| *Niphargus zagrebensis* | NA059 | KY617339 | EF617295 | KR905789 |
| *Pseudoniphargus gorbeanus* | PN0167 | NA | KY441101 | KY441043 |
| *Pseudoniphargus portosancti* | PN0131 | NA | KY441102 | MH592136 |
