## Supplementary material for "Polyphyly of the *Niphargus stygius* species group (Crustacea, Amphipoda, Niphargidae) in the Southern Limestone Alps": Tab. S3

Table S3 - List of primers used for amplification and sequencing and PCR amplification conditions.

| Marker | Direction | Purpose | Sequence | PCR conditions | Primers reference |
|---|---|---|---|---|---|
| COI | Forward | PCR + sequencing | 5'-CHACWAAYCATAAAGATATYGG-3' | 3min at 94°C, 30x (20sec at 94°C, 45sec at 50°C, 1min at 72°C), 2min at 72°C | Astrin and Stüben (2008) |
| COI | Reverse | PCR + sequencing | 5'-AWACTTCVGGRTGVCCAAARAATCA-3' | | Astrin and Stüben (2008) |
| 28S | Forward | PCR + sequencing | 5′-CAAGTACCGTGAGGGAAAGTT-3′ | 3min at 94°C, 30x (20sec at 94°C, 45sec at 45°C, 1min at 72°C), 2min at 72°C | Verovnik et al. (2005) |
| 28S | Reverse | PCR + sequencing | 5′-AGGGAAACTTCGGAGGGAACC-3′ | | Verovnik et al. (2005) |
| 28S | Forward | sequencing | 5′-AAACACGGGCCAAGGAGTAT-3′ | | Flot et al. (2010) |
| 28S | Reverse | sequencing | 5′-TATACTCCTTGGCCCGTGTT-3′ | | Flot et al. (2010) |
| ITS | Forward | PCR + sequencing | 5′-TCCGAACTGGTGCACTTAGA-3′ | 1min at 94°C, 40x (30sec at 94°C, 30sec at 53°C, 3 min at 72°C) | Flot et al. (2010) |
| ITS | Reverse | PCR + sequencing | 5′-TCCAAGCTCCATTGGCTTAT-3′ | | Flot et al. (2010) |
| ITS | Forward | sequencing | 5′-CGCTGCCATTCTCACACTTA-3′ | | Flot et al. (2010) |
| ITS | Reverse | sequencing | 5′-ACTCTGAGCGGTGGATCACT-3′ | | Flot et al. (2010) |
| ITS | Forward | sequencing | 5′-AAGGCTATAGCTGGCGATCA-3′ | | Flot et al. (2010) |
| ITS | Reverse | sequencing | 5′-TCAGCGGGTAACCTCTCCTA-3′ | | Flot et al. (2010) |
