## Supplementary material for "Polyphyly of the *Niphargus stygius* species group (Crustacea, Amphipoda, Niphargidae) in the Southern Limestone Alps": Tab. S4

Table S4 - Results of the molecular species delimitation analysis using ASAP, PTP, and the patristic distance threshold methods.

| **ASAP-ITS: tot 8** | **SP** | **ASAP-COI: tot 19** | **SP** | **Threshold-COI tot 11** | **SP** | **PTP-ITS: tot 10** | **SP** | **PTP COI: tot 23** | **SP** |
|---|---|---|---|---|---|---|---|---|---|
| FS_18.086-*N*. cf. *julius* Eisen#a | 1 | FS_12.043-*N*. cf. *julius* Mainarda | 1 | FS_12.043-*N*. cf. *julius* Mainarda | 1 | FS_16.036-*N*. cf. *julius* Obir | 1 | FS_12.043-*N*. cf. *julius* Mainarda | 1 |
| FS_18.086-*N*. cf. *julius* Eisen#b | 1 | FS_13.087-*N*. cf. *julius* Mainarda2 | 1 | FS_13.087-*N*. cf. *julius* Mainarda2 | 1 | FS_18.086-*N*. cf. *julius* Eisen#a | 1 | FS_13.087-*N*. cf. *julius* Mainarda2 | 1 |
| FS_16.036-*N*. cf. *julius* Obir | 1 | FS_13.089-*N*. cf. *julius* Pielungo | 1 | FS_13.089-*N*. cf. *julius* Pielungo | 1 | FS_18.086-*N*. cf. *julius* Eisen#b | 1 | FS_13.089-*N*. cf. *julius* Pielungo | 1 |
| FS_18.090-*N*. cf. *julius* Solimbergo | 2 | FS_18.074-*N*. cf. *julius* Solimbergo | 1 | FS_18.074-*N*. cf. *julius* Solimbergo | 1 | FS_18.074-*N*. cf. *julius* Solimbergo | 2 | FS_18.074-*N*. cf. *julius* Solimbergo | 1 |
| FS_18.074-*N*. cf. *julius* Solimbergo | 2 | FS_18.090-*N*. cf. *julius* Solimbergo | 1 | FS_18.090-*N*. cf. *julius* Solimbergo | 1 | FS_18.090-*N*. cf. *julius* Solimbergo | 2 | FS_18.090-*N*. cf. *julius* Solimbergo | 1 |
| FS_12.043-*N*. cf. *julius* Mainarda | 2 | FS_16.036-*N*. cf. *julius* Obir | 2 | FS_16.036-*N*. cf. *julius* Obir | 2 | FS_12.043-*N*. cf. *julius* Mainarda | 3 | FS_16.036-*N*. cf. *julius* Obir | 2 |
| FS_13.087-*N*. cf. *julius* Mainarda2 | 2 | FS_18.086-*N*. cf. *julius* Eisen | 2 | FS_18.086-*N*. cf. *julius* Eisen | 2 | FS_13.087-N. cf. *julius* Mainarda2 | 3 | FS_18.086-*N*. cf. *julius* Eisen | 2 |
| FS_13.089-*N*. cf. *julius* Pielungo | 2 | FS_11.097-*N*. *julius* Musi | 3 | FS_11.097-*N*. *julius* Musi | 3 | FS_13.089-*N*. cf. *julius* Pielungo | 3 | FS_11.097-*N*. *julius* Musi | 3 |
| FS_11.097-*N*. *julius* Musi | 3 | FS_11.114-*N*. *julius* Starcedat | 3 | FS_11.114-*N*. *julius* Starcedat | 3 | FS_11.097-*N*. *julius* Musi | 4 | FS_11.114-*N*. *julius* Starcedat | 3 |
| FS_11.114-*N*. *julius* Starcedat | 3 | FS_12.044-*N*. *julius* Dovica | 4 | FS_12.044-*N*. *julius* Dovica | 3 | FS_11.114-*N*. *julius* Starcedat | 4 | FS_12.044-*N*. *julius* Dovica | 4 |
| FS_12.044-*N*. *julius* Dovica#a | 3 | FS_13.097-*N*. *julius* Papipano | 4 | FS_13.097-*N*. *julius* Papipano | 3 | FS_12.044-*N*. *julius* Dovica#a | 4 | FS_14.022-*N*. *julius* Villanova | 4 |
| FS_12.044-*N*. *julius* Dovica#b | 3 | FS_14.022-*N*. *julius* Villanova | 4 | FS_14.022-*N*. *julius* Villanova | 3 | FS_12.044-*N*. *julius* Dovica#b | 4 | FS_13.097-*N*. *julius* Papipano | 5 |
| FS_13.097-*N*. *julius* Papipano | 3 | NA086-*N*. *kenki* KY617662 | 5 | NA086-*N*. *kenki* KY617662 | 4 | FS_13.097-*N*. *julius* Papipano | 4 | NA086-*N*. *kenki* KY617662 | 6 |
| FS_14.022-*N*. *julius* Villanova#a | 3 | NA087-*N*. *kenki* KY617663 | 5 | NA087-*N*. *kenki* KY617663 | 4 | FS_14.022-*N*. *julius* Villanova#a | 4 | NA087-*N*. *kenki* KY617663 | 7 |
| FS_14.022-*N*. *julius* Villanova#b | 3 | NA201-*N*. *kenki* KY617669 | 5 | NA201-*N*. *kenki* KY617669 | 4 | FS_14.022-*N*. *julius* Villanova#b | 4 | NA201-*N*. *kenki* KY617669 | 8 |
| NA086-*N*. *kenki* KY617662 | 4 | NA236-*N*. *kenki* KY617670 | 5 | NA236-*N*. *kenki* KY617670 | 4 | NA086-*N*. *kenki* KY617662 | 5 | NA236-*N*. *kenki* KY617670 | 8 |
| NA087-*N*. *kenki* KY617663 | 4 | FS_11.038-*N*. *costozzae* | 6 | FS_11.038-*N*. *costozzae* | 5 | NA087-*N*. *kenki* KY617663 | 5 | FS_11.038-*N*. *costozzae* | 9 |
| NA201-*N*. *kenki* KY617669 | 4 | FS_11.150-*N*. *costozzae* | 6 | FS_11.039-*N*. *costozzae* | 5 | NA201-*N*. *kenki* KY617669 | 6 | FS_11.039-*N*. *costozzae* | 9 |
| NA236-*N*. *kenki* KY617670 | 4 | FS_21.020-*N*. *costozzae* | 6 | FS_11.044-*N*. *costozzae* | 5 | NA236-*N*. *kenki* KY617670 | 6 | FS_11.044-*N*. *costozzae* | 9 |
| FS_11.038-*N*. *costozzae* | 5 | FS_21.022-*N*. *costozzae* | 6 | FS_11.045-*N*. *costozzae* | 5 | FS_11.038-*N*. *costozzae* | 7 | FS_11.045-*N*. *costozzae* | 9 |
| FS_11.039-*N*. *costozzae* | 5 | FS_11.039-*N*. *costozzae* | 6 | FS_11.150-*N*. *costozzae* | 5 | FS_11.039-*N*. *costozzae* | 7 | FS_11.150-*N*. *costozzae* | 9 |
| FS_11.044-*N*. *costozzae* | 5 | FS_11.044-*N*. *costozzae* | 6 | FS_11.162-*N*. *tridentinus* | 5 | FS_11.044-*N*. *costozzae* | 7 | FS_14.109-*N*. *costozzae* | 9 |
| FS_11.045-*N*. *costozzae*#a | 5 | FS_21.023-*N*. *costozzae* | 6 | FS_14.109-*N*. *costozzae* | 5 | FS_11.045-*N*. *costozzae*#a | 7 | FS_18.046-*N*. *costozzae* | 9 |
| FS_11.045-*N*. *costozzae*#b | 5 | FS_20.018-*N*. *costozzae* | 6 | FS_16.048-*N*. *costozzae* | 5 | FS_11.045-*N*. *costozzae*#b | 7 | FS_18.057-*N*. *costozzae* | 9 |
| FS_11.150-*N*. *costozzae* | 5 | FS_11.045-*N*. *costozzae* | 6 | FS_18.046-*N*. *costozzae* | 5 | FS_11.150-*N*. *costozzae* | 7 | FS_18.059-*N*. *costozzae* | 9 |
| FS_11.162-*N*. *tridentinus* | 5 | FS_14.109-*N*. *costozzae* | 6 | FS_18.057-*N*. *costozzae* | 5 | FS_11.162-*N*. *tridentinus* | 7 | FS_20.018-*N*. *costozzae* | 9 |
| FS_14.109-*N*. *costozzae* | 5 | FS_18.059-*N*. *costozzae* | 6 | FS_18.059-*N*. *costozzae* | 5 | FS_14.109-*N*. *costozzae* | 7 | FS_21.020-*N*. *costozzae* | 9 |
| FS_16.048-*N*. *costozzae* | 5 | FS_18.046-*N*. *costozzae* | 6 | FS_20.018-*N*. *costozzae* | 5 | FS_16.048-*N*. *costozzae* | 7 | FS_21.022-*N*. *costozzae* | 9 |
| FS_18.046-*N*. *costozzae* | 5 | FS_18.057-*N*. *costozzae* | 6 | FS_21.020-*N*. *costozzae* | 5 | FS_18.046-*N*. *costozzae* | 7 | FS_21.023-*N*. *costozzae* | 9 |
| FS_18.057-*N*. *costozzae* | 5 | FS_16.048-*N*. *costozzae* | 7 | FS_21.022-*N*. *costozzae* | 5 | FS_18.057-*N*. *costozzae* | 7 | FS_16.048-*N*. *costozzae* | 10 |
| FS_18.059-*N*. *costozzae* | 5 | FS_11.162-*N*. *tridentinus* | 8 | FS_21.023-*N*. *costozzae* | 5 | FS_18.059-*N*. *costozzae* | 7 | FS_11.162-*N*. *tridentinus* | 11 |
| FS_20.018-*N*. *costozzae* | 5 | FS_11.078-*N*. *brixianus* | 9 | FS_11.078-*N*. *brixianus* | 6 | FS_20.018-*N*. *costozzae* | 7 | FS_11.078-*N*. *brixianus* | 12 |
| FS_21.020-*N*. *costozzae* | 5 | FS_12.016-*N*. *brixianus* | 9 | FS_12.016-*N*. *brixianus* | 6 | FS_21.020-*N*. *costozzae* | 7 | FS_12.016-*N*. *brixianus* | 12 |
| FS_21.022-*N*. *costozzae* | 5 | FS_21.010-*N*. *brixianus* | 10 | FS_21.006-*N*. *brixianus* | 6 | FS_21.022-*N*. *costozzae* | 7 | FS_21.010-*N*. *brixianus* | 13 |
| FS_21.023-*N*. *costozzae* | 5 | FS_12.015-*N*. *brixianus* | 11 | FS_21.009-*N*. *brixianus* | 6 | FS_21.023-*N*. *costozzae* | 7 | FS_12.015-*N*. *brixianus* | 14 |
| FS_11.078-*N*. *brixianus* | 6 | FS_21.006-*N*. *brixianus* | 12 | FS_12.015-*N*. *brixianus* | 7 | FS_11.078-*N*. *brixianus* | 8 | FS_21.006-*N*. *brixianus* | 15 |
| FS_11.102-*N*. cf. *forelii* | 6 | FS_21.009-*N*. *brixianus* | 12 | FS_21.010-*N*. *brixianus* | 7 | FS_11.102-*N*. cf. *forelii* | 8 | FS_21.009-*N*. *brixianus* | 15 |
| FS_11.103-*N*. *lessiniensis* | 6 | FS_13.014-*N*. *brixianus* | 13 | FS_13.014-*N*. *brixianus* | 8 | FS_11.103-*N*. *lessiniensis* | 8 | FS_13.014-*N*. *brixianus* | 16 |
| FS_11.104-*N*. cf. *lessiniensis* | 6 | FS_11.102-*N*. cf. *forelii* | 14 | FS_11.102-*N*. cf. *forelii* | 9 | FS_11.104-*N*. cf. *lessiniensis* | 8 | FS_11.102-*N*. cf. *forelii* | 17 |
| FS_12.015-*N*. *brixianus* | 6 | FS_18.092-*N*. cf. *forelii* | 14 | FS_11.104-*N*. cf. *lessiniensis* | 9 | FS_12.015-*N*. *brixianus* | 8 | FS_18.092-*N*. cf. *forelii* | 18 |
| FS_12.016-*N*. *brixianus* | 6 | FS_11.104-*N*. cf. *lessiniensis* | 15 | FS_18.092-*N*. cf. *forelii* | 9 | FS_12.016-*N*. *brixianus* | 8 | FS_11.104-*N*. cf. *lessiniensis* | 19 |
| FS_13.014-*N*. *brixianus* | 6 | FS_11.103-*N*. *lessiniensis* | 16 | FS_11.103-*N*. *lessiniensis* | 10 | FS_13.014-*N*. *brixianus* | 8 | FS_11.103-*N*. *lessiniensis* | 20 |
| FS_18.092-*N*. cf. *forelii* #a | 6 | FS_11.091-*N*. *montellianus* | 17 | FS_11.091-*N*. *montellianus* | 11 | FS_18.092-*N*. cf. *forelii* #a | 8 | FS_11.091-*N*. *montellianus* | 21 |
| FS_18.092-*N*. cf. *forelii* #b | 6 | FS_11.096-*N*. *montellianus* | 17 | FS_11.095-*N*. *montellianus* | 11 | FS_18.092-*N*. cf. *forelii* #b | 8 | FS_11.095-*N*. *montellianus* | 21 |
| FS_21.006-*N*. *brixianus* | 6 | FS_11.095-*N*. *montellianus* | 17 | FS_11.096-*N*. *montellianus* | 11 | FS_21.006-*N*. *brixianus* | 8 | FS_11.096-*N*. *montellianus* | 21 |
| FS_21.009-*N*. *brixianus* | 6 | FS_11.092-*N*. cf. *montellianus* | 18 | FS_11.092-*N*. cf. *montellianus* | 11 | FS_21.009-*N*. *brixianus* | 8 | FS_11.092-*N*. cf. *montellianus* | 22 |
| FS_21.010-*N*. *brixianus* | 6 | FS_12.017-*N*. cf. *montellianus* | 18 | FS_12.017-*N*. cf. *montellianus* | 11 | FS_21.010-*N*. *brixianus* | 8 | FS_12.017-*N*. cf. *montellianus* | 22 |
| FS_11.091-*N*. *montellianus* | 7 | FS_12.041-*N*. cf. *montellianus* | 19 | FS_12.041-*N*. cf. *montellianus* | 11 | FS_11.091-*N*. *montellianus* | 9 | FS_12.041-*N*. cf. *montellianus* | 23 |
| FS_11.095-*N*. *montellianus* | 7 | FS_14.082-*N*. cf. *montellianus* | 19 | FS_12.045-*N*. cf. *montellianus* | 11 | FS_11.095-*N*. *montellianus* | 9 | FS_12.045-*N*. cf. *montellianus* | 23 |
| FS_11.096-*N*. *montellianus* | 7 | FS_14.110-*N*. cf. *montellianus* | 19 | FS_12.047-*N*. cf. *montellianus* | 11 | FS_11.096-*N*. *montellianus* | 9 | FS_12.047-*N*. cf. *montellianus* | 23 |
| FS_11.092-*N*. cf. *montellianus* | 8 | FS_12.045-*N*. cf. *montellianus* | 19 | FS_13.088-*N*. cf. *montellianus* | 11 | FS_11.092-*N*. cf. *montellianus* | 10 | FS_13.088-*N*. cf. *montellianus* | 23 |
| FS_12.017-*N*. cf. *montellianus* | 8 | FS_13.088-*N*. cf. *montellianus* | 19 | FS_14.082-*N*. cf. *montellianus* | 11 | FS_12.017-*N*. cf. *montellianus* | 10 | FS_14.082-*N*. cf. *montellianus* | 23 |
| FS_12.041-*N*. cf. *montellianus* | 8 | FS_14.119-*N*. cf. *montellianus* | 19 | FS_14.110-*N*. cf. *montellianus* | 11 | FS_12.041-*N*. cf. *montellianus* | 10 | FS_14.110-*N*. cf. *montellianus* | 23 |
| FS_12.045-*N*. cf. *montellianus* | 8 | FS_12.047-*N*. cf. *montellianus* | 19 | FS_14.119-*N*. cf. *montellianus* | 11 | FS_12.045-*N*. cf. *montellianus* | 10 | FS_14.119-*N*. cf. *montellianus* | 23 |
| FS_12.047-*N*. cf. *montellianus* | 8 | FS_14.214-*N*. cf. *montellianus* | 19 | FS_14.214-*N*. cf. *montellianus* | 11 | FS_12.047-*N*. cf. *montellianus* | 10 | FS_14.214-*N*. cf. *montellianus* | 23 |
| FS_13.088-*N*. cf. *montellianus* | 8 | | | | | FS_13.088-*N*. cf. *montellianus* | 10 | | |
| FS_14.082-*N*. cf. *montellianus* | 8 | | | | | FS_14.082-*N*. cf. *montellianus* | 10 | | |
| FS_14.110-*N*. cf. *montellianus* | 8 | | | | | FS_14.110-*N*. cf. *montellianus* | 10 | | |
| FS_14.119-*N*. cf. *montellianus* | 8 | | | | | FS_14.119-*N*. cf. *montellianus* | 10 | | |
| FS_14.214-*N*. cf. *montellianus* | 8 | | | | | FS_14.214-*N*. cf. *montellianus* | 10 | | |
